# The molecular basis for Pompe disease revealed by the structure of human acid α-glucosidase

**DOI:** 10.1101/212837

**Authors:** Derrick Deming, Karen Lee, Tracey McSherry, Ronnie R. Wei, Tim Edmunds, Scott C. Garman

## Abstract

Pompe disease results from a defect in human acid α-glucosidase (GAA), a lysosomal enzyme that cleaves terminal α1-4 and α1-6 glucose from glycogen. In Pompe disease (also known as Glycogen Storage Disorder type II), the accumulation of undegraded glycogen in lysosomes leads to cellular dysfunction, primarily in muscle and heart tissues. Pompe disease is an active candidate of clinical research, with pharmacological chaperone therapy tested and enzyme replacement therapy approved. Despite production of large amounts of recombinant GAA annually, the structure of GAA has not been reported until now. Here, we describe the first structure of GAA, at 1.7Å resolution. Three structures of GAA complexes reveal the molecular basis for the hundreds of mutations that lead to Pompe disease and for pharmacological chaperoning in the protein. The GAA structure reveals a surprising second sugar-binding site 34Å from the active site, suggesting a possible mechanism for processing of large glycogen substrates. Overall, the structure will assist in the design of next-generation treatments for Pompe disease.

## Introduction

The lysosomal storage disorder Pompe disease (also known as Glycogen Storage Disorder type II) is caused by loss of activity in the acid α-glucosidase (GAA) enzyme (Fukuda et al., 2007, Lim et al., 2015, Raben et al., 2002). GAA is an exo-hydrolase cleaving both α1-4 and α1-6 glucose linkages in glycogen, releasing α-glucose as a product (Fig. 1A). In the absence of functional GAA, glycogen accumulates in the lysosomes, leading to cellular dysfunction and the onset of Pompe disease symptoms, including muscle atrophy, liver enlargement, and heart defects (Hirschhorn and Reuser, 2001). The age of onset of Pompe disease correlates with GAA activity, where infantile, juvenile, and adult onset forms have an average of 0%, 5%, and 8% residual GAA activity respectively (Hirschhorn and Reuser, 2001).

**Figure 1.**
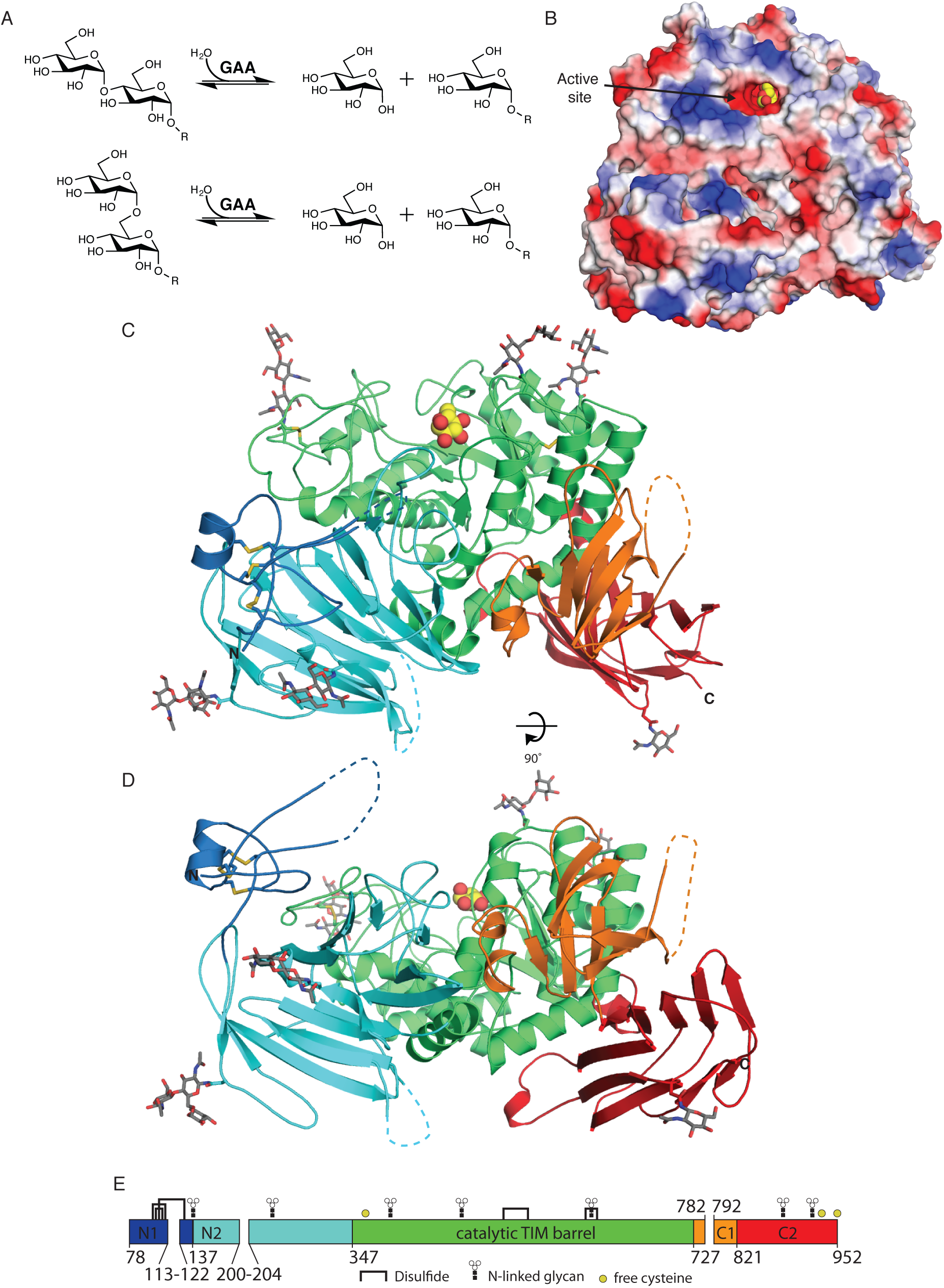
Overall Structure of GAA. (A) GAA cleaves terminal α1-4- and α1-6-linked glucose from glycogen polymers. (B) Electrostatic surface potential with the reaction product α-glucose (yellow) bound to the active site. (C-D) Ribbon diagram of GAA colored by domain, with N-linked glycans shown in grey and glucose (yellow) bound to the active site. Dotted lines indicate residues missing in electron density. (E) Schematic of the GAA polypeptide colored by domain showing the location of disulfide bonds, N-linked glycans and free cysteines. Gaps represent residues cleaved during maturation.

Pompe disease can be treated by enzyme replacement therapy, by infusion of recombinant GAA, an approach approved by the FDA in 2006 (Lim et al., 2014). Because of the number of cells requiring enzyme and inefficient uptake of recombinant enzyme, massive doses of purified recombinant GAA (up to 20 mg/kg biweekly) are required to treat Pompe disease patients, making enzyme replacement therapy expensive (Schoser et al., 2008). Additionally, large doses can provoke immune responses in Pompe disease patients (Bigger et al., 2015, Doerfler et al., 2016). Thus, GAA remains an active research target, and alternative approaches are in preclinical and clinical development. For example, current approaches include modification of the glycans on GAA to improve its uptake into the lysosome (Zhu et al., 2009, Zhu et al., 2005), addition of insulin-like growth factor II tags to improve uptake (Maga et al., 2013), gene therapy using adeno-associated-virus vectors (Doerfler et al., 2016), pharmacological chaperoning with small-molecule stabilizers, and combination therapy using both enzyme replacement and pharmacological chaperoning approaches (Khanna et al., 2014).

Pharmacological chaperones bind to the active site of their targets and thus stabilize the folded state of the protein (Fan et al., 1999). For Pompe disease, the iminosugar glucose analog 1-deoxynojirimycin (DNJ) can increase GAA activity, improving lysosomal localization of some GAA variants (Flanagan et al., 2009). Co-administration of pharmacological chaperone with recombinant GAA enzyme can increase enzyme activity in the lysosome (Khanna et al., 2012, Khanna et al., 2014). While DNJ can chaperone GAA, it also inhibits multiple glucosidases, potentially leading to unwanted effects. The identification of more specific chaperones has been hindered by a lack of structural information on GAA.

GAA is synthesized as a 952-residue zymogen and undergoes at least nine proteolytic processing steps during maturation (Hoefsloot et al., 1988, Moreland et al., 2005, Wisselaar et al., 1993). After cleavage of the 27-residue signal sequence and glycosylation in the ER, the 110 kDa GAA protein (residues 28-952) is trafficked to the Golgi, where glycans are further modified and proteolytic cleavage yields residues 57-952 (Moreland et al., 2005). Multiple cleavage events in the late endosome and lysosome produce the mature protein, containing four polypeptides with approximate masses of 3,10, 70, and 19 kDa corresponding to residues 78-113, 122-200, 204-782, and 792-952 respectively (Moreland et al., 2005) (Fig. 1E). The mature enzyme shows improved kinetics against glycogen (with a 10-fold lower K_M_ and a 2-fold higher k_cat_), but the molecular basis of the effect is not well understood (Bijvoet et al., 1998, Wisselaar et al., 1993).

To better understand the molecular basis of Pompe disease, we determined the three-dimensional structure of the mature human GAA by X-ray crystallography. Additionally, to reveal substrate recognition and pharmacological chaperoning in GAA, we determined complexes of GAA with the catalytic product glucose, the pharmacological chaperone DNJ, a disaccharide, and a trisaccharide. The structures reveal a surprising second sugar-binding site in GAA distal from the active site, suggesting a mechanism for processive catalysis of glycogen polymers. The structure allows for modeling of the locations of Pompe disease variants onto the structure, indicating that the disease is primarily a protein folding disorder, because mutations tend to disrupt the hydrophobic core of the protein.

## Results and Discussion

### Structure determination

Extensive crystallization trials with the intact GAA (residues 70-952) resulted in crystals that diffracted to 10Å resolution at best. To reduce conformational heterogeneity, we developed an *in vitro* limited protease digestion to mimic the proteolytic maturation *in vivo* and to remove loops hindering crystallization. Limited proteolysis cleaved the protein into fragments, including a doublet of 70-80 kDa and a fragment of approximately 20 kDa (Fig. 1 supplement A). The fragments migrate as a single species during size exclusion chromatography, indicating they remain associated post cleavage. Crystallization of the protease-treated GAA selected for the more highly processed fragments (Fig. 1 supplement A), leading to crystals that diffracted to 1.7Å resolution (Table 1).

**Table 1:** Crystallographic data.

| PDB Code | 5KZR | 5KZW | 5KZX |
| --- | --- | --- | --- |
| Ligand Soak | DNJ | isomaltose | maltotriose |
| Active site ligand | DNJ | glucose | glucose |
| Second site ligand | - | isomaltose | maltotriose |
| cell lengths, Å | 96.6, 102.9, 128.7 | 97.1, 102.6, 128.6 | 97.1, 102.6, 128.5 |
| Resolution, Å* | 50-1.7 (1.73-1.7) | 50-2.0 (2.04-2.0) | 50-1.9 (1.93-1.9) |
| Observations* | 870,867 (41,948) | 326,654 (21,225) | 658,306 (17,008) |
| Unique observations* | 141,597 (6,895) | 73,011 (4,969) | 90,718 (2,389) |
| Completeness, %* | 100.0 (99.9) | 84.0 (93.3) | 89.2 (47.8) |
| Multiplicity* | 6.2 (6.1) | 4.5 (4.3) | 7.3 (7.1) |
| R<sub>sym</sub>, %* | 7.6 (47.2) | 7.3 (36.7) | 8.7 (34.8) |
| <I/σ<sub>i</sub>>* | 11.8 (3.0) | 13.7 (3.5) | 12.9 (5.9) |
| R<sub>work</sub> / R<sub>free</sub>, % | 13.7 / 15.6 | 15.4 / 18.5 | 15.5 / 17.9 |
| Atoms | 7,828 | 7,803 | 7,827 |
| Protein | 6,734 | 6,742 | 6,726 |
| Carbohydrate | 87 | 218 | 229 |
| Water | 836 | 792 | 816 |
| Other | 171 | 51 | 56 |
| Average B-factor (Å<sup>2</sup>) | 22.0 | 26.1 | 17.8 |
| Protein | 19.6 | 23.9 | 14.9 |
| Carbohydrate | 49.9 | 57.0 | 51.6 |
| Water | 33.1 | 36.2 | 32.1 |
| Ramachandran plot, % |  |  |  |
| Favored | 97.6 | 97.3 | 97.4 |
| Allowed | 2.4 | 2.7 | 2.6 |
| Outlier | 0 | 0 | 0 |
| RMS deviations |  |  |  |
| Bonds, Å | 0.012 | 0.010 | 0.009 |
| Angles, ° | 1.55 | 1.47 | 1.41 |

\*Highest resolution shell appears in parentheses

### Overall GAA structure

The human GAA structure contains five domains, seven N-linked glycans, five disulfide bonds, three unpaired cysteines, and three missing loops (consistent with their removal by protease). The five domains include an N-terminal P-type trefoil domain (N1), a β-sandwich domain (N2), a (β/α)_8_ barrel containing the catalytic site, a proximal β-sandwich domain (C1), and a distal C-terminal β-sandwich domain (C2) (Fig. 1 and Fig. 1 supplement B). The N1 domain spanning residues 80-136 contains little secondary structure other than a short α-helix (100-106) and is maintained with three disulfide bonds. The N2 domain spanning residues 137-349 has a β-sandwich topology with 16 antiparallel β-strands in two β-sheets. The third domain encompasses residues 347-726 and has the characteristic (β/α)_8_ topology found in the catalytic domain of family 31 glycoside hydrolases. C-terminal to the catalytic (β/α)_8_ domain, the GAA structure contains two additional β-sandwich domains. Domain C1 spans residues 727-820 and is composed of seven antiparallel β-strands in 5- and 2-stranded β-sheets. The C2 domain has a β-sandwich topology and is composed of 10 β-strands in two 5-stranded β-sheets, with two small α-helices inserted (Fig. 1 and Fig. 1 supplement B).

The GAA structure reveals extensive post-translational modifications of the polypeptide. The N-linked glycosylation sites show clear electron density at six of the seven reported sites. The 13 cysteine residues form five disulfide bonds, three in domain N1 (82-109, 92-108, and 103-127) and two in the catalytic domain (533-558 and 647-655). The disulfide arrangement in N1 matches the 1-5, 2-4 and 3-6 connectivity found in other P-type trefoil domains (Sim et al., 2008, Wright et al., 1997). Three remaining cysteines are unpaired, including C374 near the active site and the surface-exposed C938 and C952 in the C2 domain (Fig. 1).

Proteolytic treatment prior to crystallization mimics proteolytic maturation in the late endosome and lysosome. Electron density is missing for loops 116-122, 199-205, and 782-792. Presumably, the flexible loops in the intact protein limited the resolution of the intact GAA crystals. After cleavage of the three loops, the remaining fragments remain associated by non-covalent interactions and a disulfide bond in N1 and by non-covalent interactions in domains N2 and C1. Extensive surface area remains buried in the mature GAA: 1105 Å^2^ (and the C103-C127 disulfide bond) between the 80-115 fragment and the rest of the protein; 2944 Å^2^ between the 123-200 fragment and the rest of the protein; and 2901 Å^2^ between the 793-952 fragment and the rest of the protein (Fig. 1 supplement C). Because of the large surface area buried between proteolytic fragments, the GAA subunits remain associated after proteolytic maturation (Moreland et al., 2005).

### Active site and ligand binding

The active site of GAA is an acidic pocket at the base of a conical funnel lined by hydrophobic residues in the (β/α)_8_ barrel catalytic domain (Fig. 1, Fig. 2, and Fig. 2 supplement A). The funnel and active site are formed by loops at the C-terminal ends of each of the eight β-strands forming the β-barrel. Additionally, loop 281-286 from the N2 domain contributes to substrate recognition in the active site (Fig. 2 supplement A). Thirteen residues (D282, W376, D404, L405, I441, W481, W516, D518, M519, R600, D616, W618, F649, L650, H674) define the active site within 5Å of the glucose ligand (Fig. 2).

**Figure 2.**
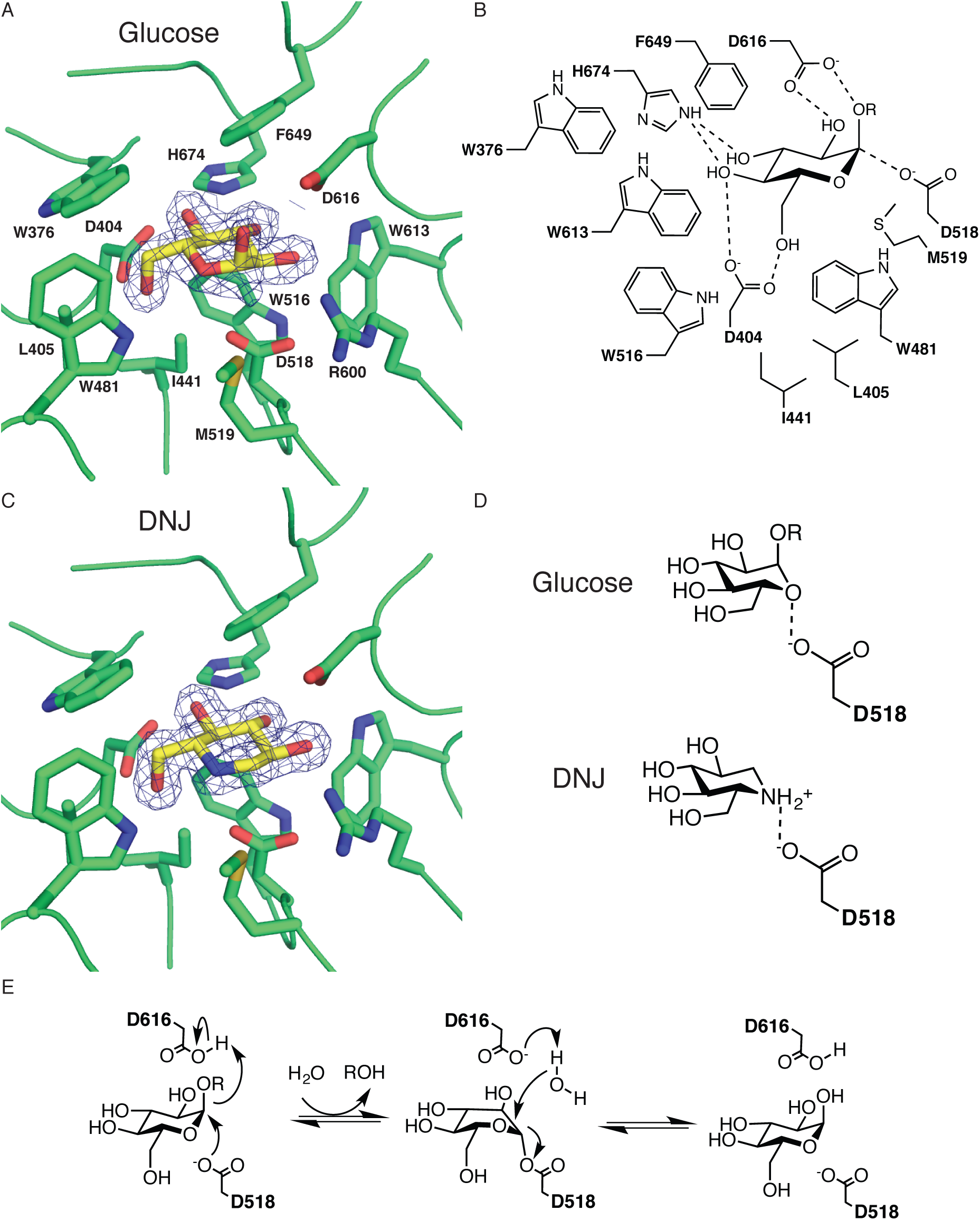
Active site and ligand binding. (A) The reaction product glucose (yellow) bound to the active site. (B) Diagram of binding interaction between GAA and glucose. (A-B) Electron density contoured at 1.0σ. (C) The pharmacological chaperone DNJ bound to the active site. (D) Comparison of the binding interaction between the heterocyclic atom of the ligand and nucleophile. (E) Proposed α-retaining reaction mechanism.

GAA uses a double-displacement reaction mechanism found in family 31 glycoside hydrolases, leading to overall retention of the anomeric configuration of the α-glucoside substrate (Bourne and Henrissat, 2001, Koshland, 1953). The nucleophile D518 falls 3.1Å from the C1 position of glucose, in position to make a nucleophilic attack on the carbon. The acid/base D616 falls on the opposite side of the cleaved glycosidic linkage, 2.9Å from the O1 of the ligand, in position to protonate the leaving group during catalysis (Fig. 2). Fitting the pH-dependence curve for GAA activity (Fig. 2 supplement B) yields two pKa values (3.6 and 5.3), which we interpret as the pKa’s for D518 and D616 respectively, consistent with the double-displacement reaction mechanism.

To better understand substrate recognition in GAA, we determined complexes of the enzyme with isomaltose and maltotriose (di- and tri-saccharide fragments of glycogen) at 2.0 and 1.9Å resolution, respectively. The structures show glucose present in the active site with no clear electron density for the remaining portion of the substrates (Fig. 2). We interpret the density as catalytic product in the active site, despite the limited GAA activity at pH 7.5 (Fig. 2 supplement B). The glucose density appears as a mixture of α and β anomers. Because of the close proximity of active site residues D282, M519, and R600, β-linked substituents beyond a hydroxyl are sterically blocked, and thus we conclude that the β-glucose appears in the active site after cleavage of α-linked substrate and subsequent ring opening of the α-glucose product. Residues D404, D616, and H674 confer glucose substrate specificity by hydrogen bonding to hydroxyls on the ligand (Fig. 2). Hydrophobic residues W376, L405, I441, W481, W516, M519, W613 and F649 contribute to active site architecture and substrate binding.

We also determined the complex of GAA bound to 1-deoxynojirimycin (DNJ) (Fig. 2), a pharmacological chaperone investigated for the treatment of Pompe disease. DNJ, a glucose analog, binds in the active site and acts as a competitive inhibitor of GAA. A global fit of substrate vs. inhibitor vs. velocity data leads to a K_i_ for DGJ of 150 +/- 8 nM (Fig. 2 supplement C).

### Second ligand binding site

Surprisingly, the maltotriose and isomaltose structures reveal a second sugar binding site, a shallow cleft in the N1 domain 34Å away from the active site (Fig. 3 and Fig. 3 supplement A). In the second site, the non-reducing ends of isomaltose and maltotriose bind identically, with hydrogen bonds between the glucose hydroxyls and the side chain of D91 and the main chain of G123, W126 and C127 (Fig. 3 and Fig. 3 supplement A). In the second site, the penultimate sugars of the two substrate fragments bind differently, where the maltotriose stacks directly over W126 and the isomaltose shifts closer to the main chain of I98. The second binding site falls close to a loop that is removed during proteolytic maturation of GAA, which might explain how glycogen affinity and enzymatic activity increase in mature GAA.

**Figure 3.**
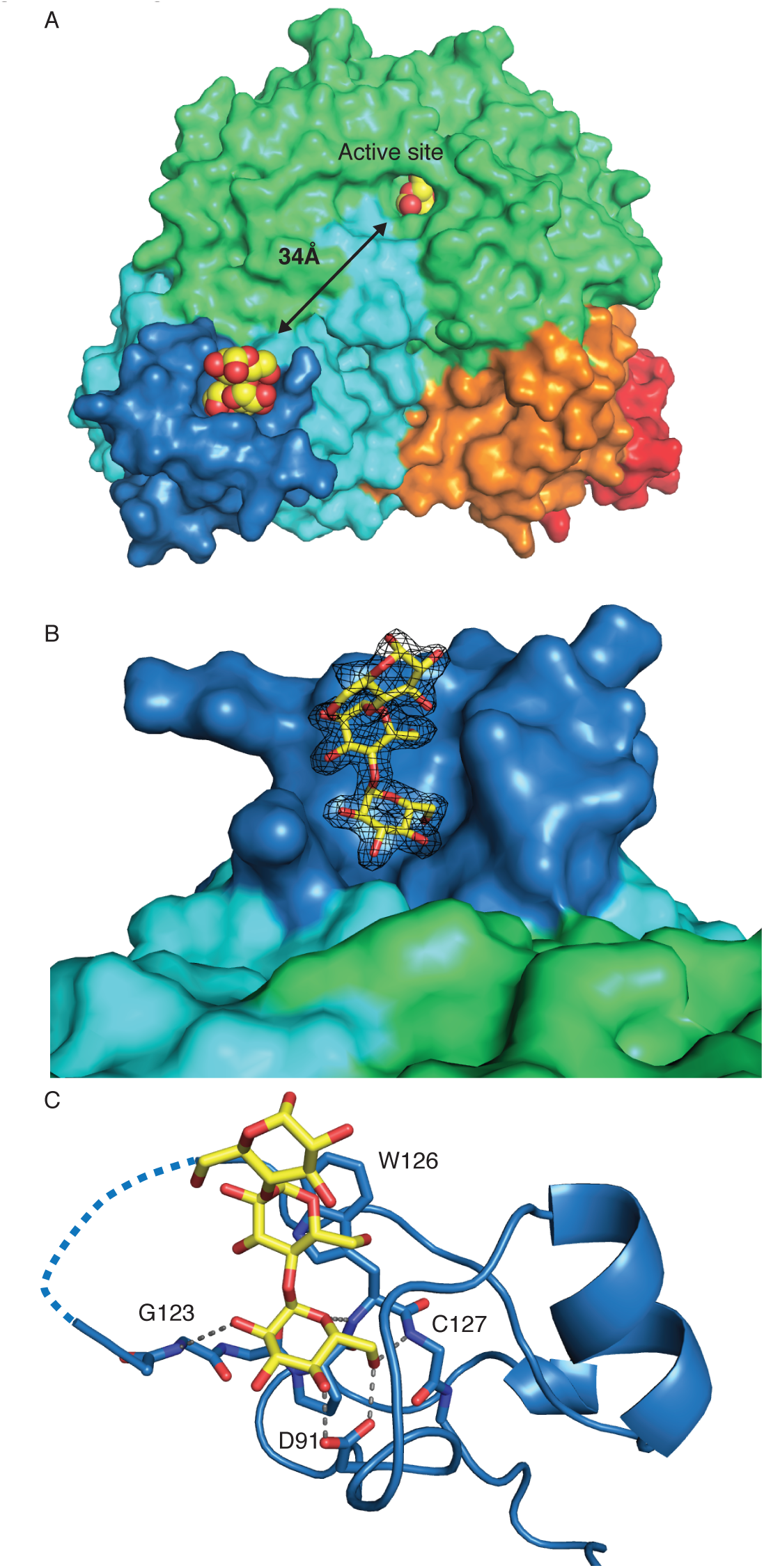
GAA contains a second substrate binding site. (A) Structure of GAA with glucose (yellow) bound to the active site and maltotriose (yellow) bound to the N-terminal domain. (B) Surface representation of the CBM in the N-terminal domain with maltotriose bound (yellow). 2Fo-Fc electron density contoured at 1.0σ. (C) Identification of key residues involved in substrate binding with polar contacts highlighted by grey dashed lines. Dotted lines represent residues missing in electron density.

Many eukaryotic family 31 glycoside hydrolases are multi-domain proteins, but the functions of the non-catalytic domains are poorly understood. The identification of a second carbohydrate-binding site in GAA suggests a functional role for the P-type trefoil domain in domain N1 as a carbohydrate-binding module (CBM). The presence of a CBM in the N-terminal domain explains the observation that the D91N variant of GAA binds carbohydrate less effectively and exhibits faster electrophoretic mobility on a starch gel (Martiniuk et al., 1990).

### Glycogen spanning the two ligand-binding sites on GAA

The second binding site offers a potential explanation of for the observation that mature GAA shows increased enzymatic activity for larger substrates. To better understand the interaction of GAA with glycogen, we modeled the GAA:glycogen interaction, guided by the two ligand-binding sites seen crystallographically (Figure 4). We computationally minimized a glycogen fragment containing 26 glucose residues with one branch point, and then manually docked the glycogen model into the GAA structure. The glycogen model fits into the two GAA binding sites with only minor adjustments of two dihedrals. The model immediately suggests a mechanism of action of GAA on glycogen, where the cleavage of the glycogen in the active site is followed by release of product from the active site coincident with glycogen binding in the second site, leading to a high effective concentration of glycogen at all stages.

**Figure 4.**
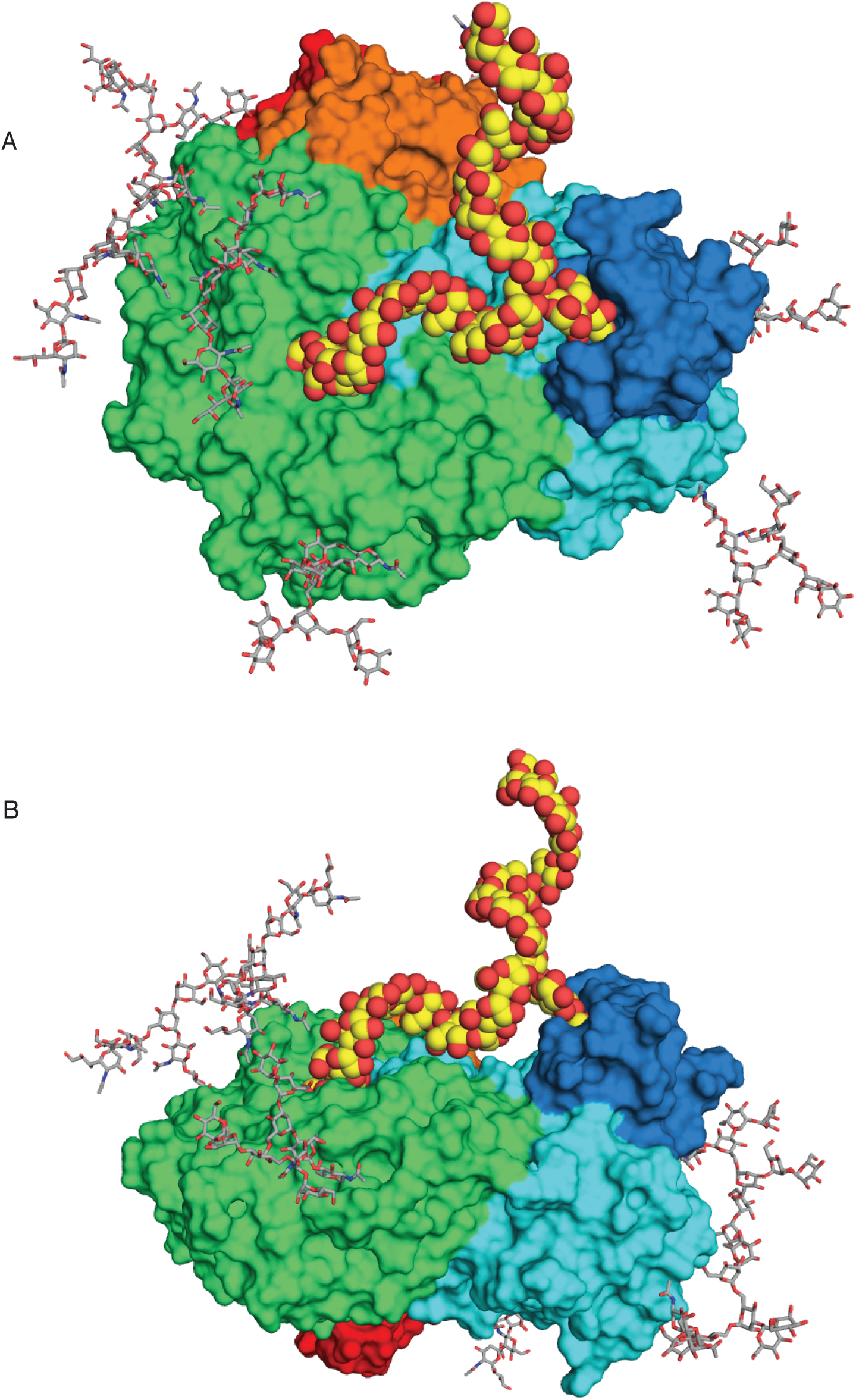
Model of GAA bound to glycogen fragment. (A) Front view of GAA bound to a glycogen fragment (yellow) showing one non-reducing end in the active site and the other non-reducing end bound the N-terminal CBM. Full length N-linked glycans shown in grey. (B) Model viewed from the back.

### Pompe disease mutations

More than 550 polymorphisms in human *GAA* have been identified (Kroos et al., 2012), and over 450 mutations are thought to be pathogenic for Pompe disease (Table 2). Of the Pompe disease-associated mutations, 245 are missense mutations leading to substitutions of single amino acids. To better understand genotype-phenotype correlations in Pompe disease, we mapped the 245 reported missense mutations onto the structure (Fig. 1 supplement B). 64% of all pathological point mutations map to the catalytic domain, 22% map to the N2 domain, and the remaining 14% map to the other three domains. Changes in buried residues are found more often than in surface residues: 23% of all buried residues have a Pompe disease-associated mutation, while only 5% of surface residues do (Fig. 4 supplement A). Because missense mutations in buried residues are most common, Pompe disease is primarily a disorder of protein misfolding.

**Table 2:** Pompe disease mutations.

| Mutation | Percent wild type activity* | Accessible surface area per side chain atom ($\text{\AA}^2$) | Mutation Category | Proposed effect | Reference |
| --- | --- | --- | --- | --- | --- |
| R11Q | 115 | ND | Signal sequence |  | (Kroos et al., 2008) |
| S46P | 100.2<sup>a</sup> | ND | Signal sequence |  | (Oba-Shinjo et al., 2009) |
| D67N | ND | ND | Pro sequence |  | (Maimaiti et al., 2009) |
| R74H | 103 | ND | ND |  | (Kroos et al., 2012) |
| R89H | 43.5 | 3.2 | Partially buried | Loss of hydrogen bonding network | (Kroos et al., 2012) |
| D91N | ND | 9.1 | Surface exposed | Mild; conservative substitution of surface-exposed residue | (Martiniuk et al., 1990) |
| C103G | 8.2<sup>a</sup> | 0 | Buried | Loss of disulfide to C127 | (Hermans et al., 2004) |
| C103R | 0 | 0 | Buried | Loss of disulfide to C127 | (Kroos et al., 2012) |
| C108S | ND | 0 | Buried | Loss of disulfide to C92 | (Liu et al., 2014) |
| C127F | 1.5 | 0.3 | Buried | Loss of disulfide to C103 | (Kroos et al., 2012) |
| L141M | ND | 11.3 | Surface exposed | Less room for larger Met | (Wan et al., 2008) |
| R154P | ND | 0 | Buried | Proline not favored in β-strand; loss of H-bonding | (Nascimbeni et al., 2008) |
| R168P | ND | 15.5 | Surface exposed | Proline not favored in β-strand; loss of ion pair to D170 | (Liu et al., 2013) |
| R168Q | ND | 15.5 | Surface exposed | Mild; loss of ion pair to D170 | (Liu et al., 2013) |
| L169P | ND | 0.1 | Buried | Proline not favored in β-sheet; loss of buried nonpolar interactions | (Gort et al., 2007) |
| R178C | 134 | 3.9 | Partially buried | Loss of hydrogen bonding; loss of ion pair to E174 | (Kroos et al., 2008) |
| R178H | ND | 3.9 | Partially buried | Less room for imidazole ring | (Chien et al., 2011) |
| R190H | 3.9<sup>a</sup> | 2.7 | Partially buried | Loss of hydrogen bonding network; loss of ion pair to D243 | (Kroos et al., 2008) |
| Y191C | 0.6 | 2 | Partially buried | Loss of hydrogen bonding with S315; loss of hydrophobic packing | (Pittis et al., 2008) |
| R199H | ND | ND | ND | *Not present in structure; processing affected? | (Martiniuk et al., 1990) |
| L208P | ND | 12.9 | Surface exposed | Loss of main chain H-bond | (Laforêt et al., 2000) |
| P217L | 13.3<sup>a</sup> | 7 | Partially buried | Proline favored in β-turn ; less room for Pro | (Oba-Shinjo et al., 2009) |
| G219R | ND | 0 | Buried | No room for larger side chain | (Fernandez-Hojas et al., 2002) |
| V220L | 100 | 0.5 | Buried | Less room for larger Leu | (Kroos et al., 2012) |
| V222M | 98 | 0 | Buried | Less room for larger Met | (Kroos et al., 2012) |
| H223R | ND | 5 | Partially buried | Mild | (Reuser et al., 1995) |
| R224P | 0<sup>a</sup> | 1.2 | Buried | Proline not favored in β-strand; loss of H-binding network and ion pair with E302 | (Labrousse et al., 2010) |
| R224Q | 0.7 | 1.2 | Buried | Loss of H-binding network and ion pair with E302 | (Kroos et al., 2012) |
| R224W | ND | 1.2 | Buried | No room for aromatic Trp | (Pittis et al., 2003) |
| T234K | 2.7 | 0.4 | Buried | No room for larger Lys | (Kroos et al., 2012) |
| T234R | 1.6 | 0.4 | Buried | No room for larger Arg | (Kroos et al., 2012) |
| A237V | ND | 0.1 | Buried | No room for larger Val | (Anneser et al., 2005) |
| F240S | ND | 0.5 | Buried | Loss of hydrophobic packing | (Sacconi et al., 2010) |
| A242V | 14 | 0.4 | Buried | Little room for larger Val | (Kroos et al., 2008) |
| L246R | ND | 0.1 | Buried | No room for larger Arg in hydrophobic pocket | (Spada et al., 2013) |
| L248P | 0 | 2.2 | Partially buried | Proline not favored in β-strand; loss of hydrophobic packing | (Kroos et al., 2008) |
| L248R | ND | 2.2 | Partially buried | No room for larger Arg in hydrophobic pocket | (Sacconi et al., 2010) |
| S251L | ND | 13 | Surface exposed | Mild | (Wan et al., 2008) |
| S254L | ND | 5.2 | Partially buried | Loss of H-bonding to main chain | (Wan et al., 2008) |
| G259V | 0 | 0.7 | Buried | Little room for larger Val, Gly-specific phi/psi | (Kroos et al., 2008) |
| E262K | ND | 0.1 | Buried | No room for larger Lys, loss of buried H-bond network | (Fernandez-Hojas et al., 2002) |
| P266S | 18<sup>b</sup> | 25.6 | Surface exposed | Mild | (Park et al., 2006) |
| T271A | 83.6<sup>a</sup> | 3.3 | Partially buried | Loss of hydrogen bonding with Q255 | (Labrousse et al., 2010) |
| D282H | ND | 6.5 | Partially buried | Loss of hydrogen bonding might network in active site | (Pardo et al., 2015) |
| P285R | ND | 0.3 | Buried | Little room for larger Arg | (Bodamer et al., 2002) |
| P285S | 4.8<sup>a</sup> | 0.3 | Buried | Mild | (Bali et al., 2011) |
| N290D | 71 | 0.1 | Buried | Change in buried hydrogen bonding network | (Kroos et al., 2012) |
| L291F | 0.7<sup>a</sup> | 0.2 | Buried | No room for larger Phe | (Kroos et al., 2008) |
| L291H | 2 | 0.2 | Buried | Less room for larger His | (Kroos et al., 2008) |
| L291P | 0.7<sup>a</sup> | 0.2 | Buried | Phi/psi angles outside of proline range; loss of main chain H-bond | (Kroos et al., 2008) |
| Y292C | ND | 0 | Buried | Loss of hydrophobic packing near active site | (Castro-Gago et al., 1999) |
| G293R | ND | 0 | Buried | No room for larger Arg | (Hermans et al., 2004) |
| H295Q | ND | 0 | Buried | Loss of buried H-bonding network | (Joshi et al., 2008) |
| Y298S | ND | 0 | Buried | Loss of hydrophobic packing | (Wan et al., 2008) |
| L299P | 0 | 0.4 | Buried | Proline not favored in β-strand; loss of main chain H-bond; loss of hydrophobic packing | (Kroos et al., 2008) |
| L299R | ND | 0.4 | Buried | No room for larger Arg | (Raben et al., 1995) |
| H308L | ND | 0.1 | Buried | Loss of buried hydrogen bonding network | (Laforêt et al., 2000) |
| H308P | ND | 0.1 | Buried | Proline not favored in β-strand; loss of main chain H-bond; loss of buried H-bonding network | (Hermans et al., 2004) |
| G309R | ND | 0 | Buried | No room for larger Arg | (Kroos et al., 1998) |
| V310G | ND | 0.5 | Buried | Loss of hydrophobic packing | (Kroos et al., 2012) |
| L312R | ND | 0.3 | Buried | No room for larger Arg; loss of hydrophobic packing | (Hermans et al., 2004) |
| N316I | 0 | 0.1 | Buried | Loss of hydrogen bonding network; no room for β-branched side chain | (Kroos et al., 2012) |
| M318K | 3.2<sup>a</sup> | 0.2 | Buried | Loss of buried hydrophobic interactions | (Kroos et al., 2008) |
| M318T | ND | 0.2 | Buried | Loss of buried hydrophobic interactions | (Zhong et al., 1991) |
| P324L | ND | 7.2 | Partially buried | Proline favored in β-turn | (Laforêt et al., 2000) |
| W330G | ND | 0.9 | Buried | Loss of buried hydrophobic packing | (Dou et al., 2006) |
| T333K | ND | 0.2 | Buried | No room for larger side chain | (Ding et al., 2015) |
| G334S | ND | 2.1 | Partially buried | Little room for larger side chain | (Manwaring et al., 2012) |
| G335E | 0.2 | 1.5 | Buried | No room for larger side chain; Gly-specific phi/ psi angles | (Kroos et al., 2012) |
| G335R | 0.8<sup>a</sup> | 1.5 | Buried | No room for larger side chain; Gly-specific phi/ psi angles | (Kroos et al., 2008) |
| P347R | 1 | 0.1 | Buried | N-cap of α-helix; no room for larger Arg | (Kroos et al., 2008) |
| V350M | ND | 0 | Buried | Little room for larger Met | (Quenardelle et al., 2015) |
| L355P | ND | 0.2 | Buried | Proline kinks α-helix; loss of main chain H-bond | (Hermans et al., 2004) |
| G359R | 54.3<sup>a</sup> | 10.8 | Surface exposed | Glycine specific phi/psi angles | (Müller-Felber et al., 2007) |
| P361L | 3.9<sup>a</sup> | 0.3 | Buried | Little room for Leu | (Lam et al., 2003) |
| W367R | ND | 0.5 | Buried | Loss of hydrophobic packing | (Palmer et al., 2007) |
| L369P | 0.2 | 0.2 | Buried | No room for Pro; loss of main chain H-bond | (Niño et al., 2013) |
| H372L | ND | 0 | Buried | Loss of buried H-bonding network | (Van den Hout et al., 2004) |
| L373R | ND | 0.2 | Buried | No room for larger side chain | (Bali et al., 2012) |
| C374R | ND | 0 | Buried | No room for larger Arg; disrupts active site | (Hermans et al., 2004) |
| R375H | ND | 3.4 | Partially buried | Loss of H-bonding network; disrupts active site | (Bali et al., 2011) |
| R375L | 0.3 | 3.4 | Partially buried | Loss of H-bonding network; disrupts active site | (Pittis et al., 2008) |
| G377R | 0<sup>b</sup> | 5.2 | Partially buried | Glycine specific phi/psi angles | (Kishnani et al., 2006) |
| M391V | ND | 0 | Buried | Mild; perturbs hydrophobic core | (Kroos et al., 2012) |
| P397L | 0 | 1.6 | Buried | Mild; less room for Leu | (Kroos et al., 2012) |
| Q401R | 5 | 0.1 | Buried | Little room for larger Arg | (Kroos et al., 2008) |
| W402R | ND | 0 | Buried | Loss of hydrophobic packing | (Hoefsloot et al., 1990) |
| N403K | ND | 0 | Buried | No room for larger Lys; loss of H-bonding network | (GAA Database Consortium, 2016) |
| D404G | ND | 2 | Partially buried | Loss of substrate recognition | (Nilsson et al., 2014) |
| D404N | ND | 2 | Partially buried | Loss of Hydrogen bond acceptor, might affect substrate recognition | (Amartino et al., 2006) |
| L405P | ND | 1.1 | Buried | Disrupts hydrogen bonding network in active site; perturbs active site geometry | (Hermans et al., 2004) |
| M408V | ND | 0 | Buried | little room for β-branched side chain | (Fernandez-Hojas et al., 2002) |
| S410F | 111 | 25.3 | Surface exposed | Mild | (Kroos et al., 2008) |
| D413E | ND | 0.4 | Buried | No room for larger Glu | (Dlamini et al., 2008) |
| D419V | 21.3 | 27.5 | Surface exposed | Mild | (Kroos et al., 2012) |
| M427T | ND | 0.6 | Buried | Loss of hydrophobic packing | (Ding et al., 2015) |
| Q429R | ND | 18.2 | Surface exposed | Mild | (Chien et al., 2015) |
| Q433K | ND | 22.3 | Surface exposed | Mild | (Ravaglia et al., 2012) |
| R437C | ND | 6.6 | Partially buried | loss of H-bonding network; loss of hydrophobic packing | (Lam et al., 2003) |
| R437H | ND | 6.6 | Partially buried | loss of H-bonding network; less room for aromatic His | (Turaça et al., 2015) |
| M439K | ND | 0 | Buried | loss of hydrophobic packing | (Park et al., 2006) |
| V442M | ND | 0.9 | Buried | Less room for larger Met | (Chien et al., 2011) |
| P444R | ND | 0.8 | Buried | No room for larger Arg | (Remiche et al., 2014) |
| A445P | ND | 2.6 | Partially buried | Mild; less room for Pro | (Montalvo et al., 2006) |
| Y455C | ND | 1.3 | Buried | Loss of hydrophobic packing; loss of main chain H-bond | (Gort et al., 2007) |
| Y455F | ND | 1.3 | Buried | Loss of main chain H-bond | (Hermans et al., 2004) |
| P457H | 0 | 2.7 | Partially buried | Little room for larger His | (Kroos et al., 2012) |
| P457L | 10 | 2.7 | Partially buried | Little room for larger Leu | (Kroos et al., 2008) |
| Y458C | 45 | 4.2 | Partially buried | Loss of hydrophobic packing | (Kroos et al., 2012) |
| D459N | ND | 7.4 | Partially buried | Mild; some H-bonding effect | (Wan et al., 2008) |
| G461S | 0 | 0 | Buried | No room for larger side chain | (Kroos et al., 2008) |
| L462P | ND | 17.1 | Surface exposed | Loss of main chain H-bond; Pro causes kink in α-helix | (Hu et al., 2015) |
| V466F | 0 | 0 | Buried | No room for larger Phe | (Kroos et al., 2008) |
| V466G | ND | 0 | Buried | Loss of hydrophobic packing | (Palmer et al., 2007) |
| G478R | ND | 0 | Buried | No room for larger Arg | (Reuser et al., 1995) |
| K479N | ND | 14.6 | Surface exposed | Mild | (Wan et al., 2008) |
| W481R | ND | 6 | Partially buried | Loss of hydrophobic packing | (58) |
| P482L | ND | 1.7 | Buried | Proline favored in β-turn; no room for larger Leu | (Gort et al., 2007) |
| P482R | 0<sup>b</sup> | 1.7 | Buried | Proline favored in β-turn; no room for larger Arg | (Kroos et al., 2008) |
| G483R | 0 | 20.7 | Surface exposed | Glycine-specific phi/psi angles | (Kroos et al., 2008) |
| G483V | 0<sup>a</sup> | 20.7 | Surface exposed | Glycine-specific phi/psi angles | (Kroos et al., 2008) |
| A486P | 0<sup>a</sup> | 0 | Buried | Little room for larger Pro; loss of main chain H-bond | (Oba-Shinjo et al., 2009) |
| F487S | 0 | 0.1 | Buried | Loss of hydrophobic packing | (Kroos et al., 2008) |
| D489N | ND | 1.3 | Buried | Loss of H-bond network, loss of main chain H-bond | (Montalvo et al., 2006) |
| D489Y | ND | 1.3 | Buried | No room for larger Tyr; loss of H-bond network | (Fu et al., 2014) |
| D489G | ND | 1.3 | Buried | loss of H-bond network | (Bali et al., 2012) |
| P493L | ND | 21.6 | Surface exposed | Mild, Pro initiates α-helix | (Schneider et al., 2013) |
| W499R | ND | 0 | Buried | Loss of hydrophobic packing | (Joshi et al., 2008) |
| M502V | ND | 0.9 | Buried | Mild; less room for β-branched side chain | (Montalvo et al., 2006) |
| G514R | ND | 0 | Buried | No room for larger Arg | (McCready et al., 2007) |
| M515K | ND | 0.3 | Buried | Introduction of charge into hydrophobic pocket | (Fujimoto et al., 2013) |
| M519T | ND | 2.6 | Partially buried | Disrupts hydrophobic packing in active site | (Reuser et al., 1995) |
| M519V | ND | 2.6 | Partially buried | Mild active site perturbation | (Huie et al., 1994) |
| E521K | ND | 0.1 | Buried | No room for larger Lys; loss of ion pair with R600 in the active site | (Hermans et al., 1991) |
| E521Q | 0<sup>b</sup> | 0.1 | Buried | loss of ion pair w/ R600 in the active site | (Kroos et al., 2008) |
| E521V | ND | 0.1 | Buried | loss of ion pair w/ R600 in the active site | (Liu et al., 2013) |
| P522A | 0 | 0 | Buried | Loss of hydrophobic packing | (Müller-Felber et al., 2007) |
| P522S | 0.8 | 0 | Buried | Loss of hydrophobic packing | (Kroos et al., 2008) |
| P522T | ND | 0 | Buried | Loss of hydrophobic packing | (McCready et al., 2007) |
| S523Y | 0 | 1.4 | Buried | Little room for larger Tyr; near active site | (Kroos et al., 2012) |
| F525Y | 13.4<sup>a</sup> | 8.6 | Partially buried | Less room for larger Tyr; perturbs active site | (Labrousse et al., 2010) |
| S529V | ND | 2.3 | Partially buried | Loss of main chain H-bond; less room for β-branched side chain | (Tsunoda et al., 1996) |
| P545L | ND | 0 | Buried | No room for larger Leu | (Hermans et al., 1994) |
| V548F | ND | 13.2 | Surface exposed | Mild; some loss of hydrophobic packing | (Case et al., 2008) |
| G549R | ND | 21.7 | Surface exposed | Mild; glycine specific phi/psi angles | (Hermans et al., 2004) |
| L552P | ND | 0.2 | Buried | Loss of hydrophobic packing | (Bodamer et al., 2002) |
| T556A | ND | 0.6 | Buried | Mild; some loss of H-bonding | (Loureiro Neves et al., 2013) |
| I557F | 2.7<sup>a</sup> | 0 | Buried | No room for larger Phe | (Labrousse et al., 2010) |
| C558S | 2.1<sup>a</sup> | 3.2 | Partially buried | Loss of disulfide with C533 | (Alcántara-Ortigoza et al., 2010) |
| S566P | ND | 6.5 | Partially buried | Phi/psi angle outside of proline range | (Hermans et al., 1994) |
| H568L | ND | 0 | Buried | loss of H-bonding network | (Reuser et al., 1995) |
| H568Q | ND | 0 | Buried | Mild; some changes in H-bonding network | (Alejandre et al., 2012) |
| N570K | 0.9<sup>a</sup> | 5.1 | Partially buried | Loss of some hydrogen bonding | (Kroos et al., 2008) |
| H572Q | 0<sup>a</sup> | 0 | Buried | Mild; loss of pi stacking with W279 | (Kroos et al., 2008) |
| N573H | 0 | 0.4 | Buried | Mild; less room for larger His | (Kroos et al., 2008) |
| N573K | ND | 0.4 | Buried | Little room for larger Lys | (Bali et al., 2012) |
| Y575C | 2 | 0.1 | Buried | Loss of hydrophobic packing | (Kroos et al., 2012) |
| Y575S | ND | 0.1 | Buried | Loss of hydrophobic packing | (Hermans et al., 2004) |
| G576D | 0 | 0 | Buried | No room for larger Asp | (Kroos et al., 2008) |
| G576R | 3.8 | 0 | Buried | No room for larger Arg | (Kroos et al., 2012) |
| G576S | ND | 0 | Buried | No room for larger Ser | (Reuser et al., 1995) |
| E579K | ND | 0 | Buried | Less room for larger Lys | (Hermans et al., 2004) |
| S583F | 0 | 0 | Buried | No room for larger Phe; loss of H-bond to W499 | (Kroos et al., 2008) |
| R585K | 94 | 12.9 | Surface exposed | Mild; conservative substitution of surface-exposed residue | (Kroos et al., 2008) |
| R585M | ND | 12.9 | Surface exposed | Mild | (Laforêt et al., 2000) |
| R591W | ND | 2.8 | Partially buried | Loss of ion pair to D513 | (Montagnese et al., 2015) |
| R594H | 0.1 | 2.1 | Partially buried | Loss of ion pair to D860; loss of hydrogen bonding network | (Kroos et al., 2012) |
| R594P | 1.2<sup>a</sup> | 2.1 | Partially buried | Loss of ion pair to D860; loss of hydrogen bonding network | (Kroos et al., 2008) |
| S599Y | 0 | 0 | Buried | No room for larger Tyr; loss of H-bond to N520 | (Kroos et al., 2008) |
| R600C | ND | 0.8 | Buried | Active site residue; loss of ion pair with E521; loss of H-bonding network | (Tsuji et al., 2000) |
| R600H | ND | 0.8 | Buried | Active site residue; loss of ion pair with E521; loss of H-bonding network | (Ko et al., 1999) |
| R600L | ND | 0.8 | Buried | Active site residue; loss of ion pair with E521; loss of H-bonding network | (McCreedy et al., 2007) |
| S601L | 0.4 | 0.1 | Buried | Less room for larger Leu; loss of H-bond with N280 | (Kroos et al., 2012) |
| S601W | ND | 0.1 | Buried | No room for larger Trp; loss of H-bond with N280 | (Palmer et al., 2007) |
| T602A | 5.8 | 0 | Buried | Mild; loss of some hydrophobic packing | (Kroos et al., 2012) |
| G605D | ND | 4.7 | Partially buried | No room for larger Asp; Gly-specific phi/psi | (Muraoka et al., 2011) |
| G607D | ND | 0.3 | Buried | No room for larger Asp | (Hermans et al., 2004) |
| A610V | 5<sup>b</sup> | 0 | Buried | No room for larger Val | (Müller-Felber et al., 2007) |
| G611D | ND | 0 | Buried | No room for larger Asp | (Kroos et al., 2012) |
| H612Q | ND | 0 | Buried | Mild; loss of ion pair with E262 | (Montalvo et al., 2006) |
| H612Y | 0<sup>a</sup> | 0 | Buried | Mild; loss of ion pair with E262; loss of H-bonding network | (Oba-Shinjo et al., 2009) |
| T614K | 1.7<sup>a</sup> | 0.3 | Buried | Little room for larger Lys | (Kroos et al., 2008) |
| G615R | ND | 0 | Buried | No room for larger Arg; Gly-specific phi/psi; near active site | (Ko et al., 1999) |
| D616N | ND | 8.8 | Partially buried | Active site residue critical for mechanism | (Lin et al., 2010) |
| V617A | 65 | 1.2 | Buried | Mild; loss of hydrophobic packing | (Kroos et al., 2008) |
| S619N | ND | 0.1 | Buried | Mild; less room for larger Asn | (Kroos et al., 2006) |
| S619R | ND | 0.1 | Buried | Less room for larger Arg | (Pipo et al., 2003) |
| S627F | ND | 1.1 | Buried | No room for larger Phe | (Case et al., 2008) |
| S627P | 0.5 | 1.1 | Buried | No room for larger Pro; loss of main chain H-bond; Pro causes kink in α-helix | (Kroos et al., 2012) |
| P629L | 49 | 1.5 | Buried | Mild; Pro creates kink in α-helix | (Kroos et al., 2012) |
| N635K | 0<sup>a</sup> | 0 | Buried | No room for larger Lys;<br>loss of H-bonding network | (Oba-Shinjo et al., 2009) |
| G638V | ND | 0.5 | Buried | No room for larger Val | (Van den Hout et al., 2004) |
| G638W | ND | 0.5 | Buried | No room for larger Trp | (Joshi et al., 2008) |
| L641V | ND | 0.2 | Buried | Mild; less room for larger Leu | (Turaça et al., 2015) |
| G643R | ND | 0 | Buried | No room for larger Arg | (Hermans et al., 1993) |
| A644P | ND | 0.4 | Buried | Little room for larger Pro | (Gort et al., 2007) |
| D645E | 1.9 | 0.2 | Buried | Less room for larger Asp;<br>perturbs active site | (Hermans et al., 1993) |
| D645H | ND | 0.2 | Buried | Little room for larger Asp;<br>perturbs active site | (Lin and Shieh, 1995) |
| D645N | ND | 0.2 | Buried | Mild | (Huie et al., 1998) |
| D645Y | 0<sup>b</sup> | 0.2 | Buried | Little room for larger Asp;<br>perturbs active site | (Gort et al., 2007) |
| C647W | ND | 0 | Buried | Loss of disulfide to C658 | (Huie et al., 1994) |
| G648D | 0 | 0 | Buried | No room for larger Asp;<br>Gly-specific phi/psi | (Kroos et al., 2012) |
| G648S | ND | 0 | Buried | Little room for larger Asp;<br>Gly-specific phi/psi | (Huie et al., 1998) |
| T653N | ND | 0.5 | Buried | Less room for larger Asn;<br>perturbs 647-658 disulfide | (Chien et al., 2011) |
| S654P | ND | 22.2 | Surface exposed | Mild; less room for larger Pro;<br>loss of main chain H-bond | (Wan et al., 2008) |
| R660C | ND | 0.1 | Buried | Loss of hydrogen bonding network | (Palmer et al., 2007) |
| R660H | ND | 0.1 | Buried | Loss of hydrogen bonding network | (Pipo et al., 2003) |
| W661G | 2 | 0 | Buried | Loss of hydrophobic packing | (Kroos et al., 2008) |
| G665R | ND | 0 | Buried | No room for larger Arg | (Fu et al., 2014) |
| M671R | ND | 0 | Buried | No room for larger Arg | (Kishnani et al., 2006) |
| M671K | ND | 0 | Buried | Little room for larger Lys;<br>loss of hydrophobic packing | (Ding et al., 2015) |
| R672Q | ND | 0.1 | Buried | Loss of ion pair to D645;<br>perturbs active site | (Huie et al., 1998) |
| R672W | ND | 0.1 | Buried | Loss of ion pair to D645;<br>perturbs active site | (Huie et al., 1998) |
| Q682R | ND | 0.4 | Buried | Less room for larger Arg | (Ayca et al., 2014) |
| E689K | ND | 20 | Surface exposed | Mild | (Huie et al., 1994) |
| R702C | ND | 0 | Buried | Loss of hydrogen bonding network | (Montalvo et al., 2004) |
| R702H | 13<sup>b</sup> | 0 | Buried | Perturbs hydrogen bonding network | (Qiu et al., 2007) |
| R702L | 0.4 | 0 | Buried | Loss of hydrogen bonding network | (Kroos et al., 2012) |
| L705P | ND | 0.5 | Buried | Loss of main chain H-bond; Pro causes kink in α-helix | (Turaça et al., 2015) |
| T711R | 86<sup>a</sup> | 1.1 | Buried | little room for larger Arg | (Qiu et al., 2007) |
| L712P | 8 | 5.4 | Partially buried | Loss of main chain H-bond; Pro causes kink in α-helix | (Kroos et al., 2008) |
| V718I | 127 | 10.1 | Surface exposed | Mild; room for additional methyl group | (Kroos et al., 2012) |
| V723M | ND | 0 | Buried | Less room for larger Met | (Qiu et al., 2007) |
| A724D | ND | 0 | Buried | Less room for larger Asp in hydrophobic pocket | (Park et al., 2015) |
| R725P | ND | 0.9 | Buried | Loss of H-bonding network; loss of main chain H-bond; unfavorable phi/psi angles for Pro | (Wan et al., 2008) |
| R725W | ND | 0.9 | Buried | Loss of H-bonding network; loss of ion pair with E730; less room for aromatic Trp | (Hermans et al., 1993) |
| P726R | ND | 0 | Buried | No room for larger Arg | (Ishigaki et al., 2012) |
| T737N | 0<sup>a</sup> | 0 | Buried | Less room for larger Asn | (Kroos et al., 2008) |
| Q743K | 4.1 | 1.2 | Buried | Less room for larger Lys; loss of H-bonding network | (Kroos et al., 2012) |
| Q743R | 0.8 | 1.2 | Buried | Less room for larger Arg; loss of H-bonding network | (Kroos et al., 2008) |
| W746C | 29.4<sup>c</sup> | 0 | Buried | Loss of hydrophobic packing | (Wan et al., 2008) |
| W746G | 2.1<sup>c</sup> | 0 | Buried | Loss of hydrophobic packing | (Labrousse et al., 2010) |
| W746R | 3.5<sup>c</sup> | 0 | Buried | Loss of hydrophobic packing | (Niño et al., 2013) |
| W746S | 40.1<sup>c</sup> | 0 | Buried | Loss of hydrophobic packing | (Kroos et al., 2008) |
| G759A | ND | 28.7 | Surface exposed | Mild | (Sampaolo et al., 2013) |
| E762K | ND | 20.2 | Surface exposed | Mild | (Lin et al., 2010) |
| Y766C | ND | 0.7 | Buried | Loss of hydrophobic packing; loss of hydrogen bond with D734 | (Herzog et al., 2012) |
| P768L | ND | 0.3 | Buried | Less room for larger Leu | (Yang et al., 2014) |
| P768R | ND | 0.3 | Buried | Less room for larger Arg | (Reuser et al., 1995) |
| W772R | 5 | 0.2 | Buried | Loss of hydrophobic packing | (Kroos et al., 2008) |
| I780V | ND | 6.4 | Partially buried | Mild; loss of some hydrophobic packing | (Martiniuk et al., 1990) |
| H799D | ND | 16.9 | Surface exposed | Mild | (Montagnese et al., 2015) |
| V816I | ND | 1 | Buried | Mild; less room for extra methyl group | (Hermans et al., 1993) |
| R819P | 0.4 | 6.7 | Partially buried | Loss of main chain H-bond | (Kroos et al., 2012) |
| A880D | 0 | 0.2 | Buried | Less room for larger Asp in hydrophobic pocket | (Hermans et al., 2004) |
| L901Q | ND | 0 | Buried | loss of hydrophobic packing | (Kroos et al., 2004) |
| P913R | ND | 0.1 | Buried | No room for larger Arg | (Herzog et al., 2012) |
| V916F | 0 | 0 | Buried | Little room for larger Phe | (Kroos et al., 2012) |
| S924P | 167<sup>b</sup> | 29.2 | Surface exposed | Mild | (Kishnani et al., 2006) |
| T927I | ND | 24.4 | Surface exposed | Mild | (Hermans et al., 1993) |
| L935P | 0.5<sup>a</sup> | 0 | Buried | Loss of main chain H-bond | (Kroos et al., 2008) |
| V949D | ND | 0.1 | Buried | No room for larger Asp | (Becker et al., 1998) |

\* References for activity and identification of mutants is the same unless indicated by superscript: (a) Kroos et al. 2012, (b) Kroos et al. 2008, (c) Niño et al. 2013. ND, not determined.

## Materials and Methods

### Protease treatment of GAA

Human GAA was expressed in Chinese hamster ovary cells (Moreland et al., 2005). GAA (in 50mM sodium phosphate pH 6.2, 4% mannitol and 0.01% Tween 80) was incubated with proteinase K at a molar ratio of 280:1 at 37°C for 20 hours. The protease was removed by size exclusion chromatography on a Superdex 75 10/300 GL column (GE) equilibrated with 50mM sodium phosphate pH 6.2 and 250mM NaCl.

### X-ray crystallography

Protease-treated GAA was concentrated to 7-20mg/ml in 20mM Bis-Tris, pH 6. The protein was mixed 1:1 to 1:3 with crystallization solution containing 100mM sodium HEPES, pH 7.5, 2% (v/v) PEG 400 and 2M ammonium sulfate. Crystals took two to six months to grow in sitting drop vapor diffusion experiments. Crystals were transferred into harvest buffer (100mM sodium HEPES, pH 7.5, 3% (v/v) PEG 400, and 2.2M ammonium sulfate) and flash cooled in harvest buffer with 20% glycerol added. To form complexes, crystals were soaked in harvest buffer supplemented with ligand, either 2mM DNJ for 12 hours, 10% (w/v) isomaltose for 2-10 minutes, or 10% (w/v) maltotriose for 2-10 minutes. Prior to cooling, the ligand soaks were shifted to harvest buffer containing 2mM DNJ and 20% glycerol or harvest buffer containing 20% saccharide. X-ray diffraction data were collected on beam line 14-1 of Stanford Synchrotron Radiation Lightsource at 1.1808Å wavelength. Diffraction data were indexed and integrated using iMosflm, and scaled and reduced in space group *P*2_1_2_1_2_1_ using Aimless (Evans and Murshudov, 2013) and Ctruncate (Stein and Ballard, 2009). Molecular replacement was performed in AMoRe (Navaza, 1994) from the CCP4 suite (Cowtan et al., 2011) using the 44% identical N-terminal subunit of human maltase-glucoamylase (PDB code 2QLY) (Sim et al., 2008), resulting in an initial R-factor of 46%. Subsequent building and refinement was performed in Coot (Emsley and Cowtan, 2004) and Refmac5 (Murshudov et al., 2011), respectively. Figures were constructed in Pymol (Schrödinger, 2015) and PovScript (Fenn et al., 2003).

### Mutation analysis

Accessible surface area per residue was calculated using Areaimol in CCP4 (Cowtan et al., 2011) and then ported to Excel for further analysis. Residues with an accessible surface area per side chain atom possessing less than 2 Å^2^, between 2-9 Å^2^, and above 9 were considered buried, partially buried or exposed respectively.

### Modeling and docking of glycogen fragment

A model of branched glycogen containing 26 glucose residues molecules was energy minimized using SWEET2 (Bohne et al., 1998). Docking of the minimized fragment was performed manually and was guided by electron density at both binding sites.

### Enzyme Kinetics

Kinetic parameters were determined with and without the inhibitors DNJ and glucose. 380nM GAA was incubated with 0-11μM DNJ or 0-600mM glucose in reaction buffer (100mM sodium phosphate citric acid, pH 4.5, and 100mM NaCl) at 37°C before the addition of an equal volume of 0-33mM 4-nitrophenyl-α-D-glucopyranoside substrate in reaction buffer. Aliquots of the reaction were removed every 2.5 minutes for 12.5 minutes and diluted 30-fold into 200mM sodium borate, pH 10.2. The amount of product was determined by measuring the absorbance of *para*-nitrophenylate at 400nM using an extinction coefficient of 18.2mM^−1^cm^−1^. Enzymatic parameters K_m_, K_i_, and V_max_ were calculated in Prism Pro (GraphPad) using a global fit of the Michaelis-Menten equation for competitive inhibition:

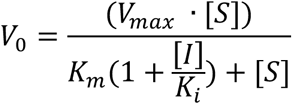

A small K_i_ for DNJ from the above global fit indicated DNJ was a tight binder, so we refit the data in Prism Pro using the Morrison equation for tight binding inhibitors:

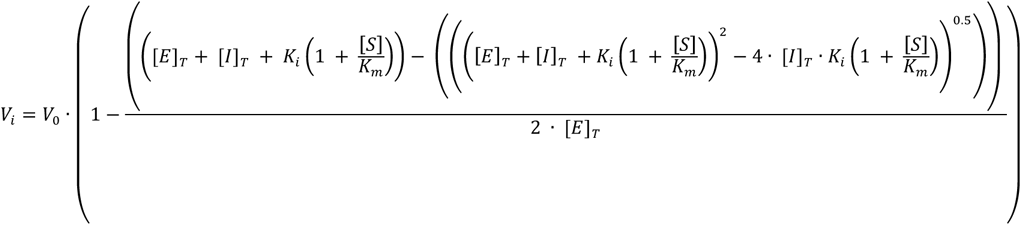

The GAA enzyme activity was measured as a function of pH, using 6.6mM 4-nitrophenyl-α-D-glucopyranoside in 100 mM sodium phosphate citric acid buffer with pH values ranging from 3-7 using 190nM GAA. Amount of product was measured as stated above. The data was fit in Prism Pro using the following equation where K_1_ and K_2_ describe the K_a_ values for the catalytic nucleophile and acid/base, respectively (Cornish-Bowden, 2004).

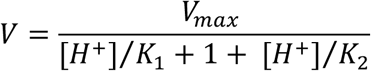

## Acknowledgements

This work was funded by NIH grant R01 DK76877 to S.C.G. and by NIH grant T32 GM008515 to D.D. We thank Matt Metcalf for prior contributions to the project, and Mike Dion for assistance. We gratefully acknowledge Vivian Stojanoff and the NSLS-SSRL user transition program supported jointly by the LSBR, NSLSII and SMB, SSRL under NIH-NIGMS grants P41GM111244 and P41GM103393, and DOE BER contracts DE-SC0012704. SSRL is operated under DOE BES contract #DE-AC02-76SF00515.

## Additional information

### Funding

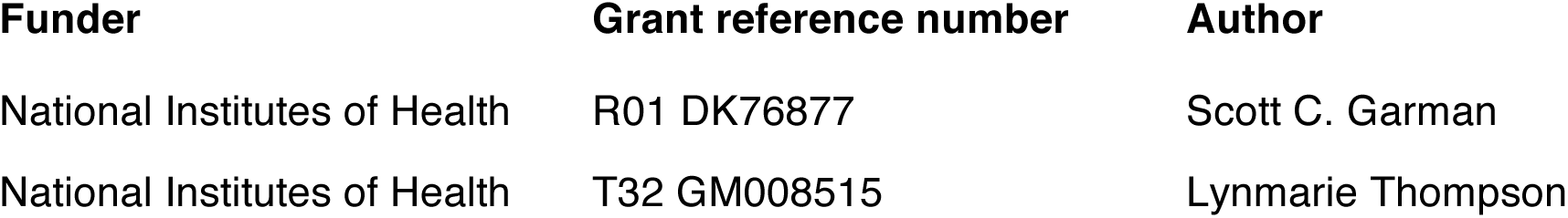

The funders had no role in study design, data collection and interpretation, or the decision to submit the work for publication.

## Author contributions

D.D. and S.C.G. designed experiments, performed experiments, analyzed data, and wrote the manuscript. K.L., T.M., R.R.W., and T.E. produced recombinant GAA and suggested experiments. All authors read and commented on the manuscript.

## Data Deposition

Atomic coordinates and structure factors have been deposited in the Protein Data Bank (www.pdb.org) under accession codes 5KZR, 5KZW, and 5KZX.

## Figure Legends

**Figure 1 Supplement A-.**
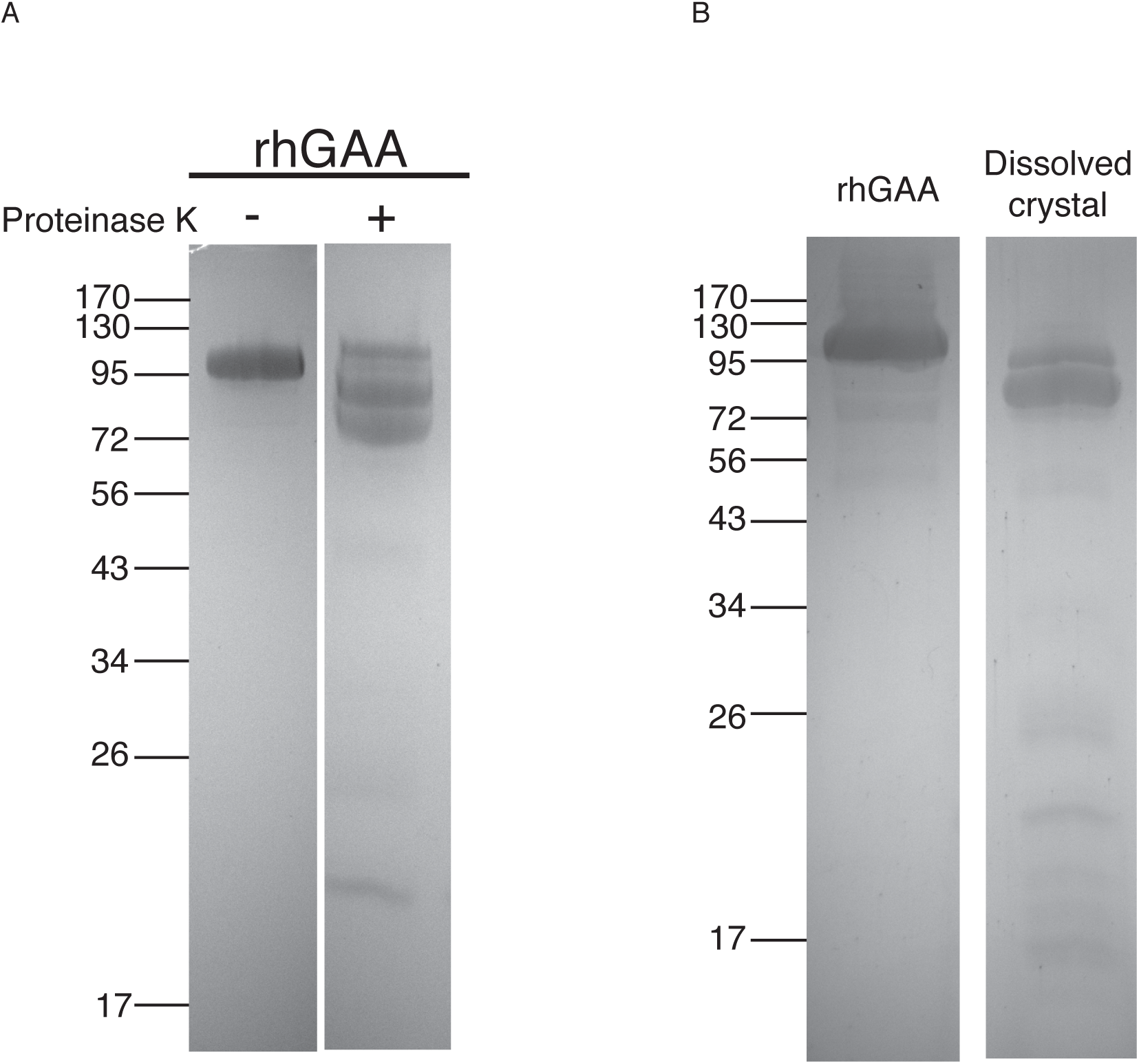
Protease treatment of rhGAA leads to crystals. (A) Reducing SDS-PAGE shows protease treatment resulted in two new bands at molecular weight 70-80 kDa and a third new band at 20 kDa. The predicted 10 and 3 kDa fragments presumably migrated off the 12% gel. (B) Reducing SDS-PAGE of dissolved crystals shows the crystallization process selected for cleaved GAA.

**Figure 1 Supplement B-.**
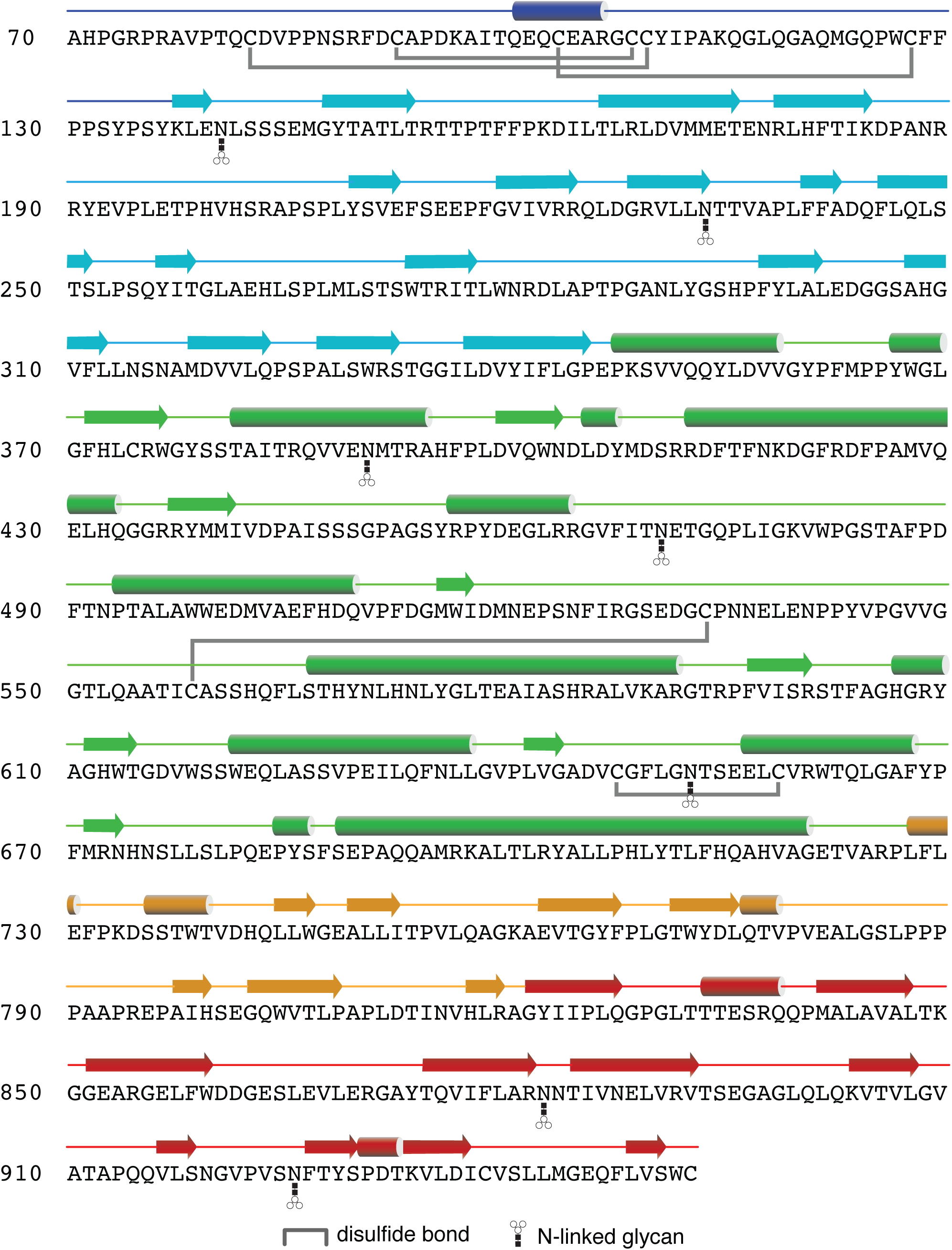
Sequence and structure of GAA. The primary sequence of human GAA is shown (colored by domain as in figure 1), along with secondary structure assignment (cylinders and arrows), disulfides (in grey) and glycans (as cartoons).

**Figure 1 Supplement C.**
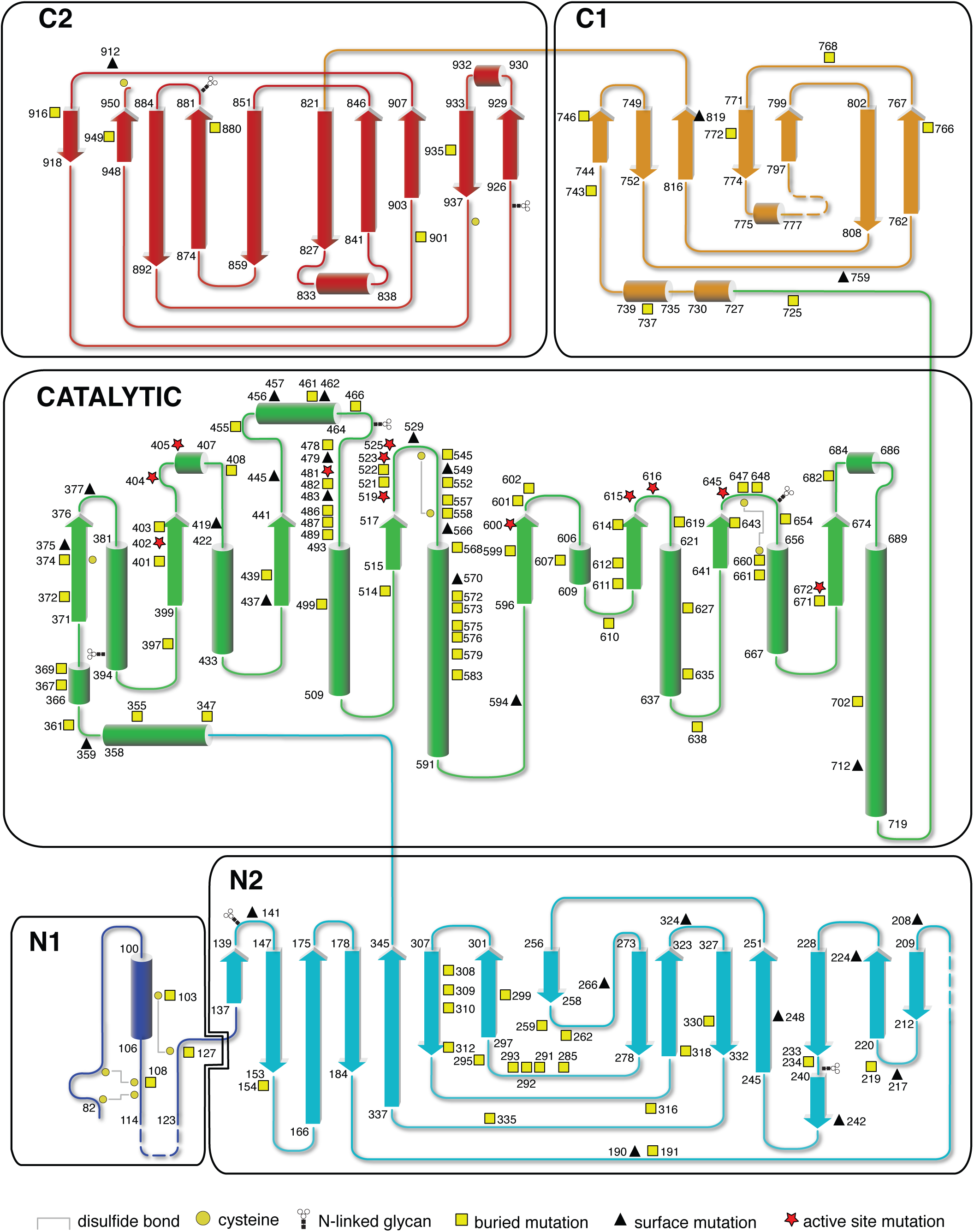
Pathological point mutations. Pathological point mutations are identified as surface exposed, buried, or active site and mapped onto the topology of GAA.

**Figure 1 Supplement D.**
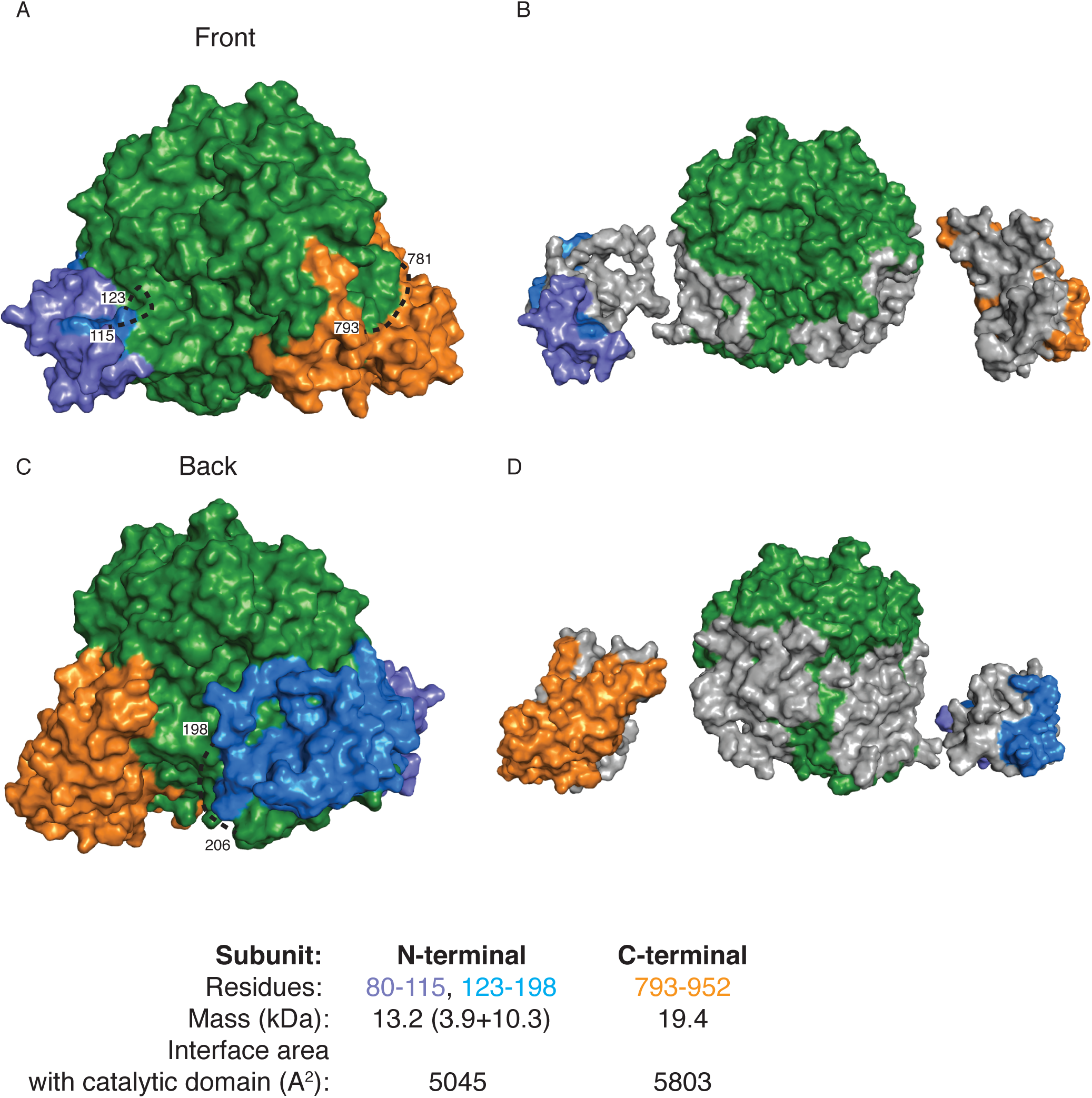
Noncovalent subunit association after proteolytic cleavage. (A,C) Surface representation of GAA colored by subunit. (B,D) Separated polypeptides with the buried interfaces colored in grey. The table shows the amino acid residues in each fragment and the buried surface area in each interface.

**Figure 2 Supplement A.**
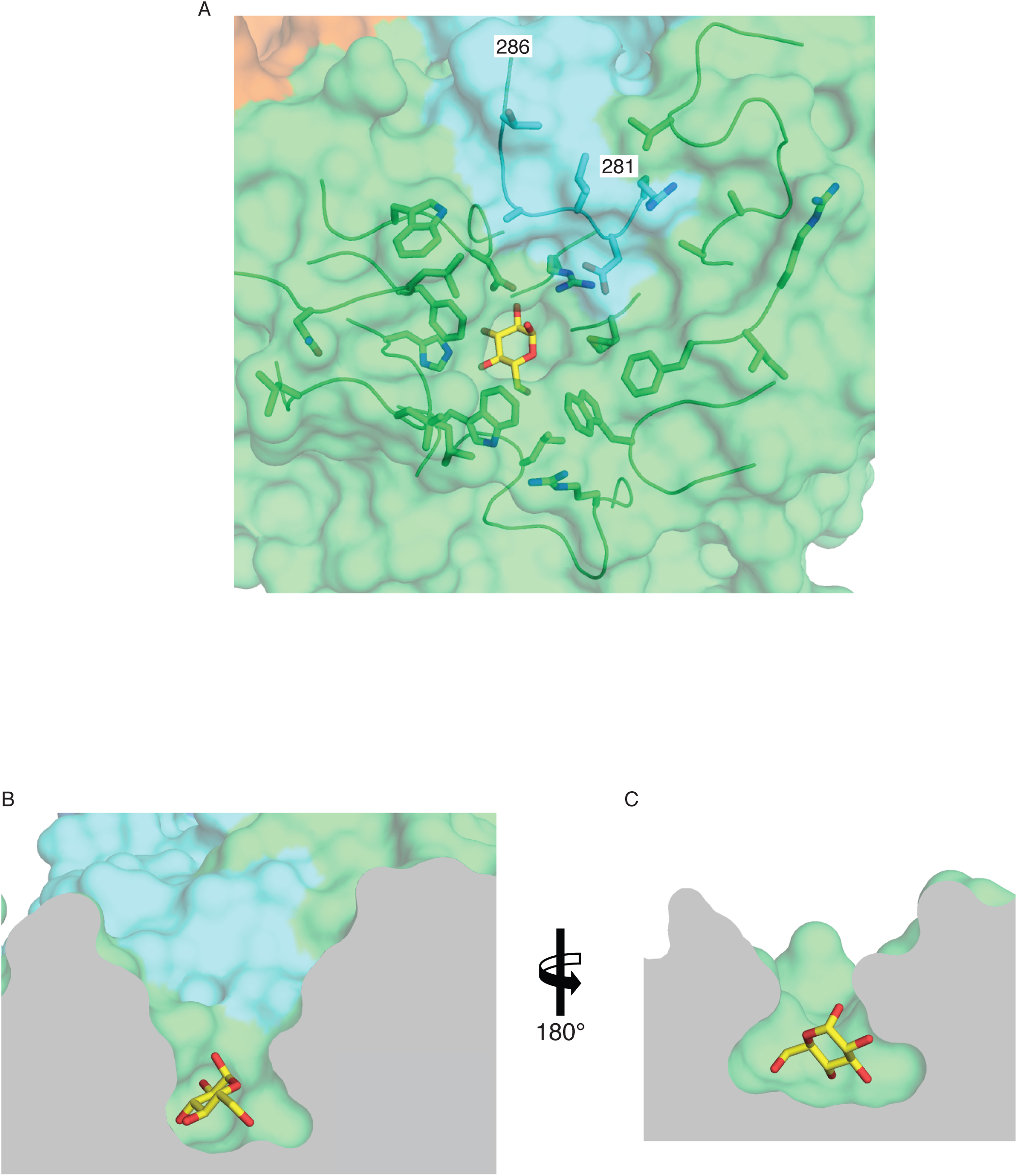
GAA contains a funnel-shaped active site. (A) Top view showing α-glucose (yellow) bound at the bottom of the active site funnel. The catalytic domain (green) and the N2 domain (cyan) are shown. A collection of hydrophobic and charged residues (shown as sticks) line the active site cavity. (B-C) Cutaway views from the sides.

**Figure 2 Supplement B.**
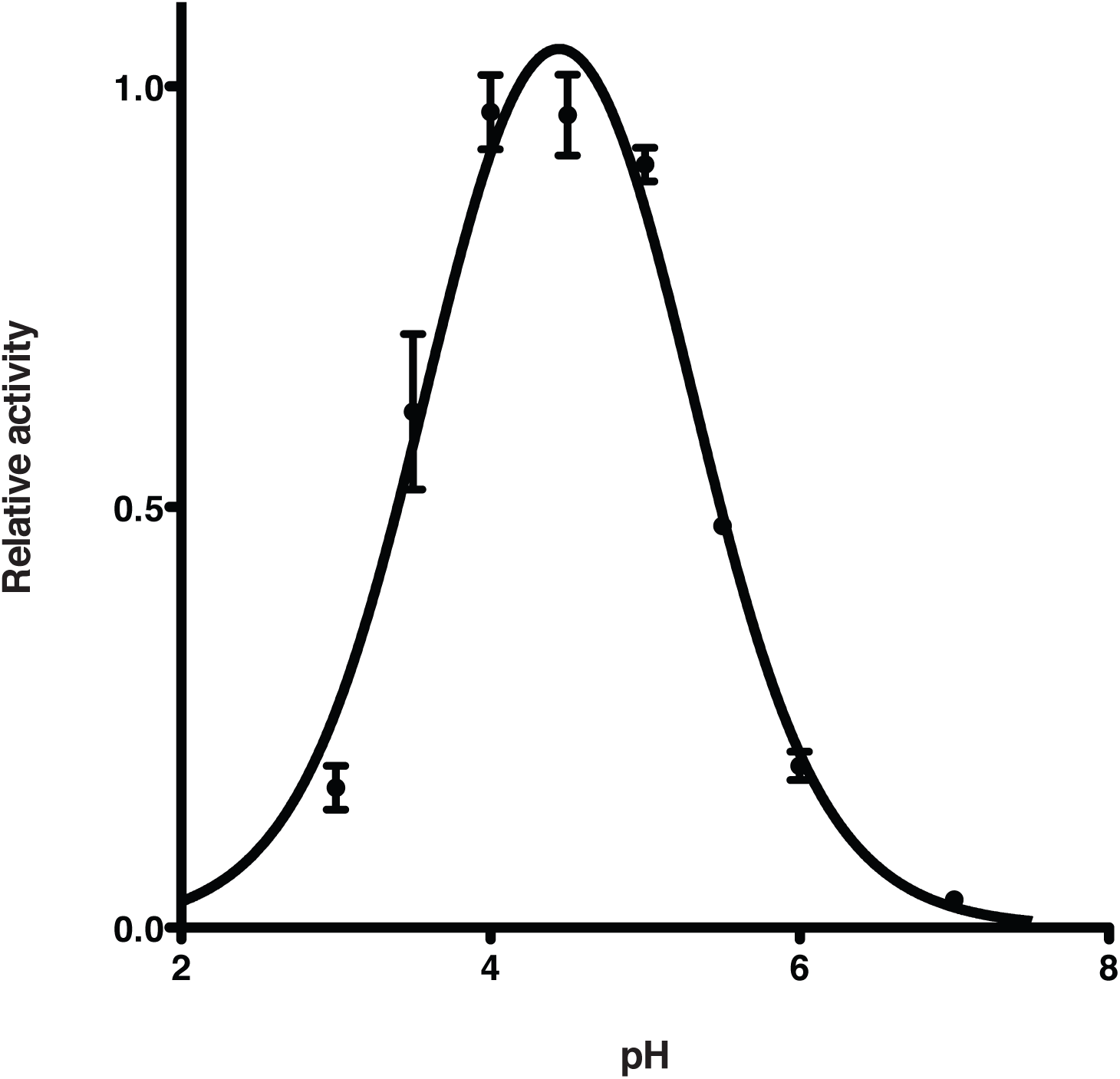
pH profile of GAA activity. Profile shows maximal activity around pH 4.5. Data were fit to equation 3, with revealing pKa values of 3.6 and 5.3 for the two ionizable groups. Error bars show the standard deviation of two replicates.

**Figure 2 Supplement C.**
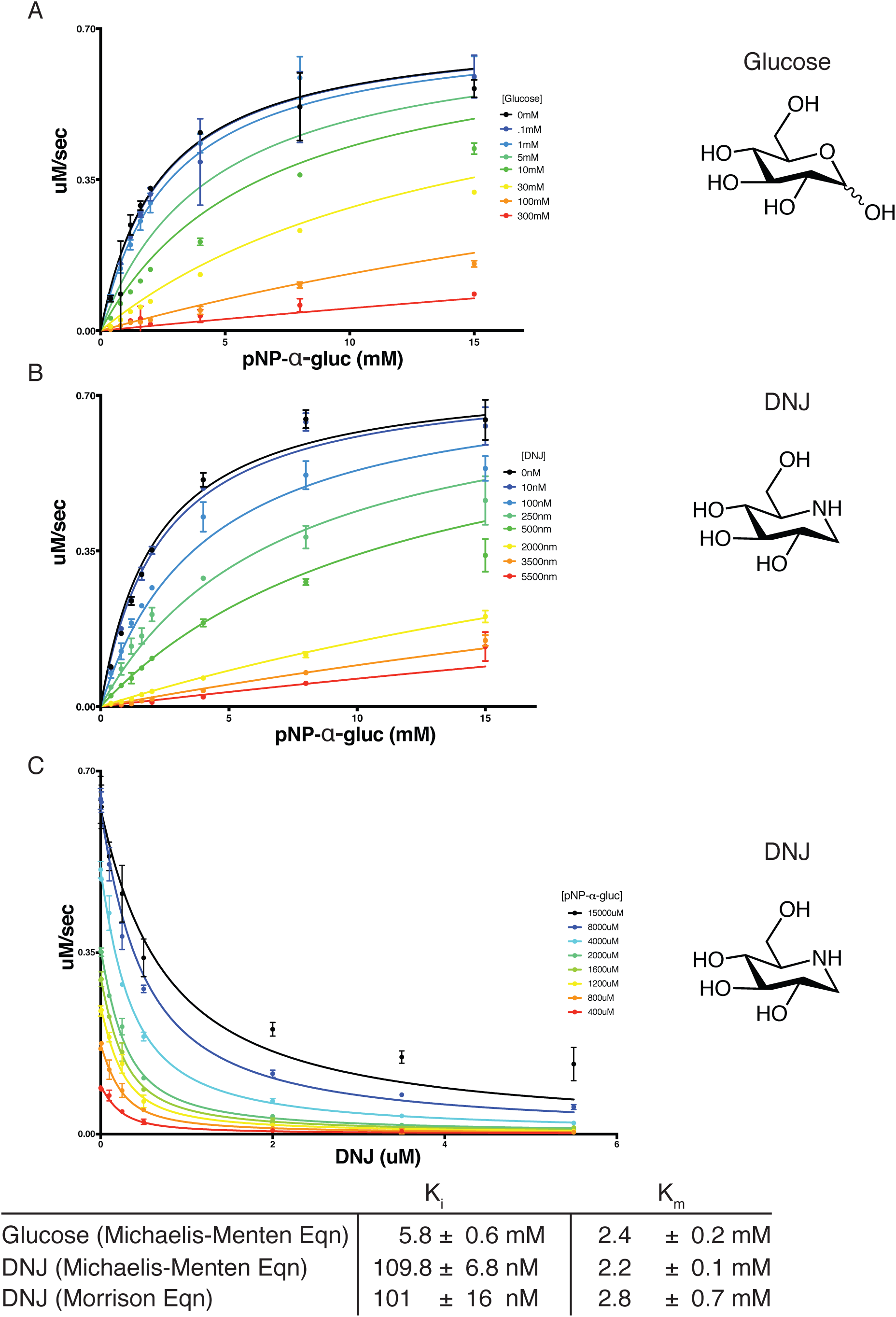
Inhibition of GAA by glucose and DNJ. (A) Inhibition of GAA by glucose, using a global fit to equation 1, the Michaelis-Menten equation for a competitive inhibitor. (B-C) Inhibition of GAA by DNJ. Data were initially fit using equation 1, then refit using the Morrison equation (Equation 2) because DNJ is a tight-binding inhibitor. Error bars show the standard deviation of two replicates.

**Figure 2 Supplement D.**
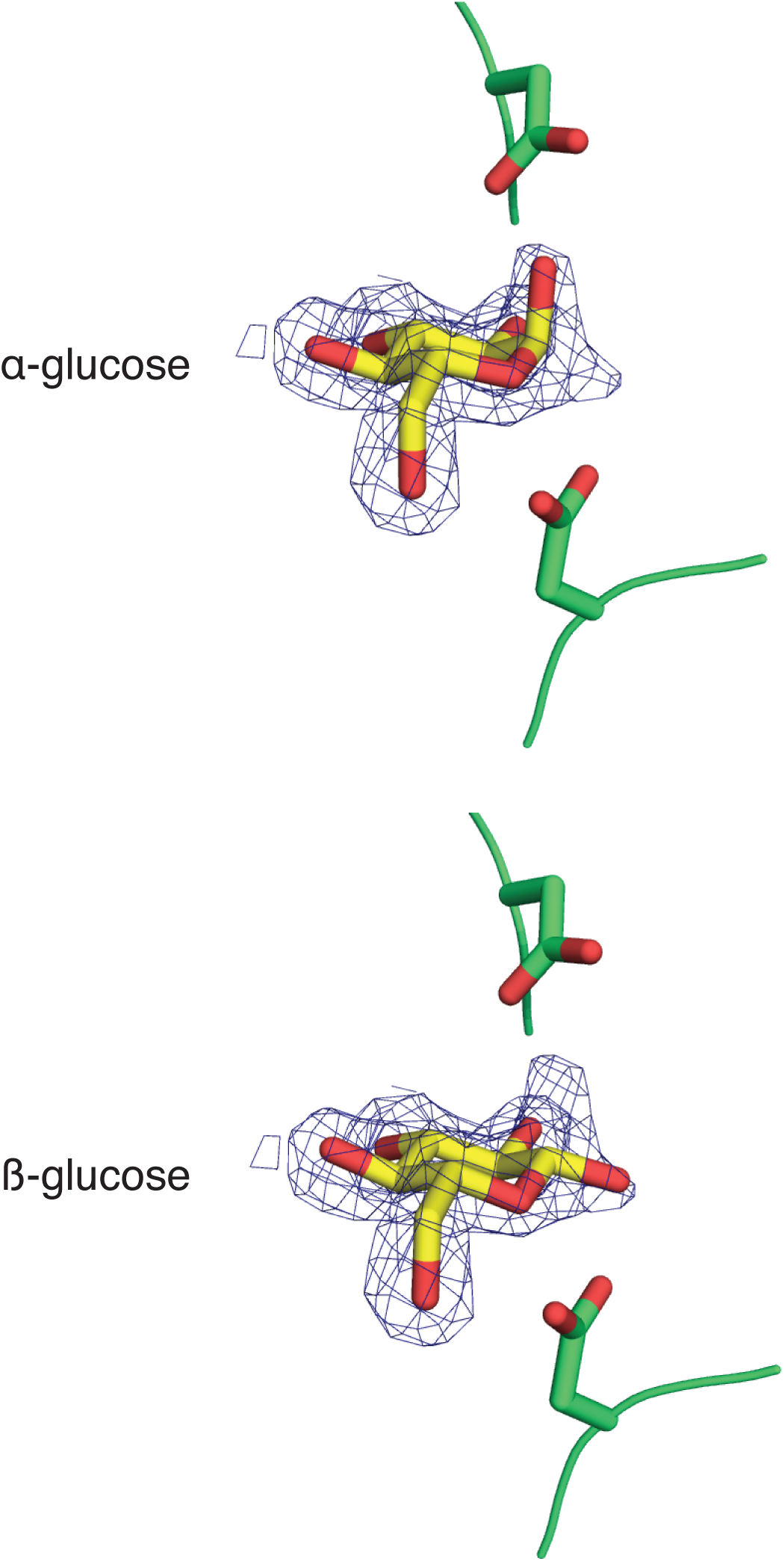
Both α and β anomers of glucose bind to the active site. Electron density reveals both α and β anomers of glucose monosaccharide bind to the active site. Larger β-linked adducts are sterically prevented from binding. 2Fo-Fc electron density is contoured at 1σ.

**Figure 3 Supplement A.**
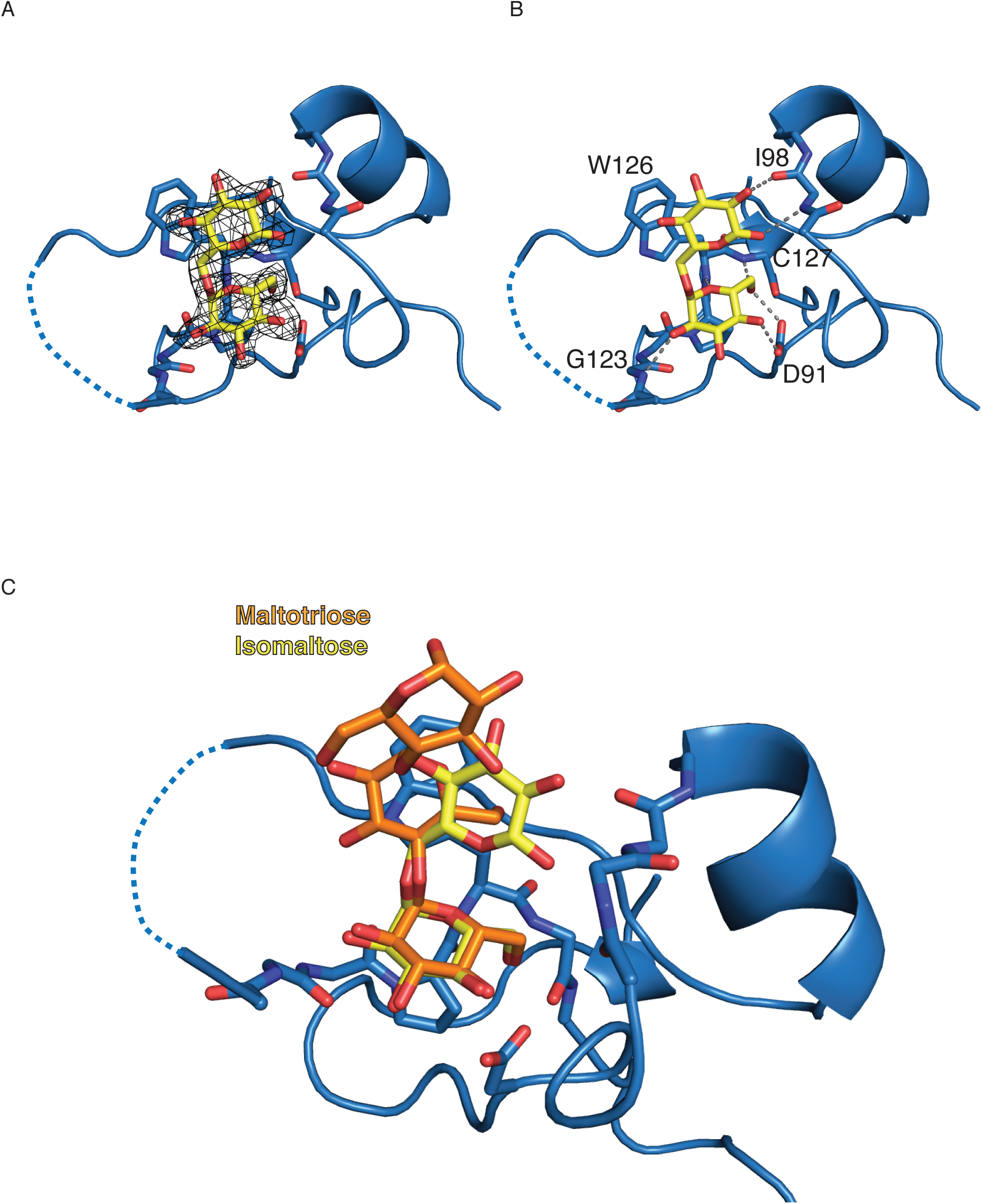
Binding of glycogen fragments to the second binding site. (A) Binding of isomaltose to the N-terminal domain with 2Fo-Fc electron density is contoured at 1σ. Dotted lines show residues absent in the electron density. (B) Key residues in ligand binding are labeled, with polar contacts highlighted by grey dashed lines. (C) Comparison of isomaltose and maltotriose bound to the second binding site.

**Figure 4 Supplement A.**
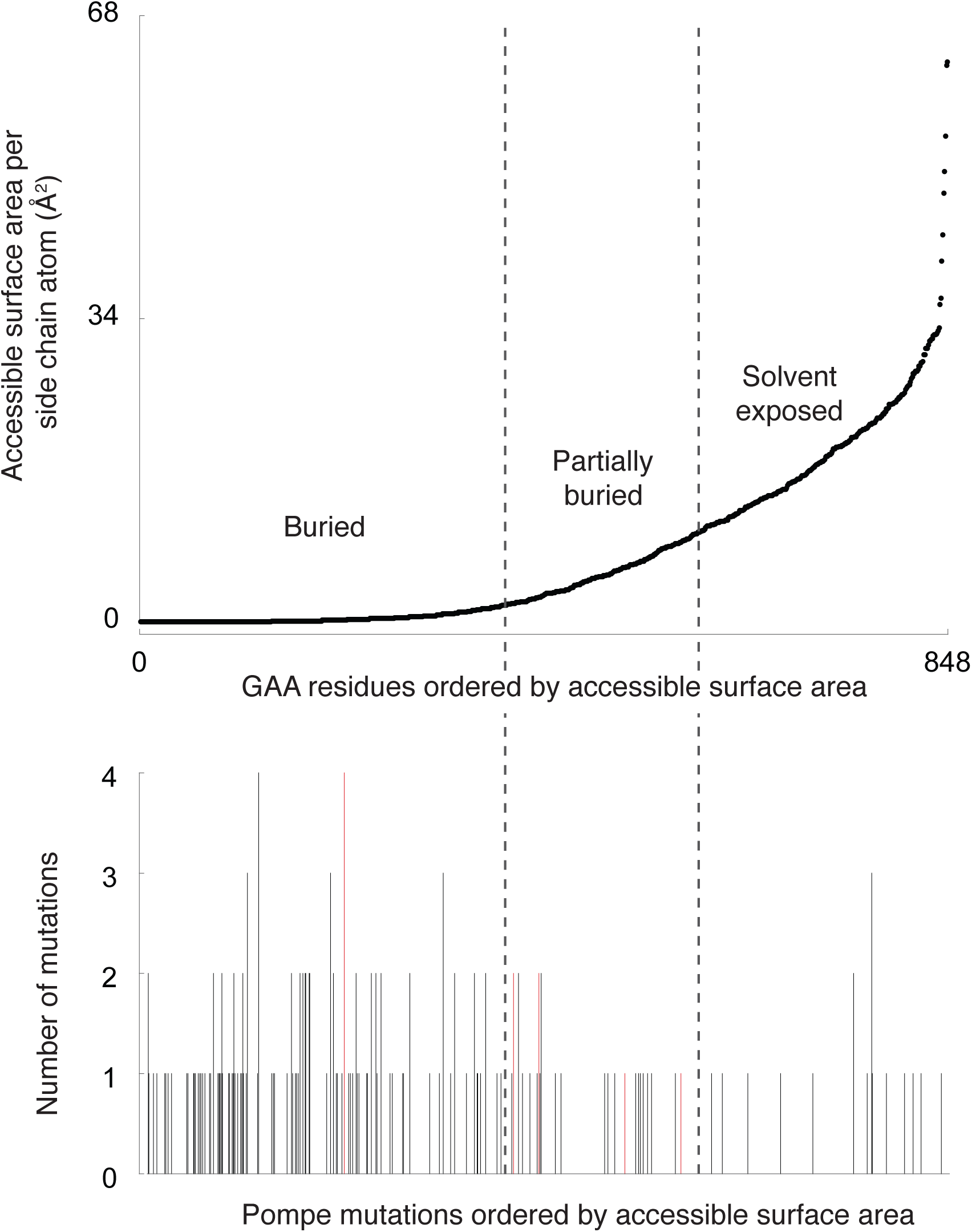
Most pathological point mutations occur in buried residues. (A) The accessible surface area per side chain atom was calculated and residues plotted in ascending order. (B) Using the same residue order as panel A, point mutations causing Pompe disease were plotted. Mutations affecting the active site are shown in red.

## Footnotes

**Conflict of Interest** D.D. declares no conflict of interest. S.C.G has received travel and speaking honoraria from Genzyme-Sanofi and Amicus. K.L., T.M., R.R.W., and T.E. are present or former employees of Sanofi.

